# Nascent Polypeptide Associated Complex–*alpha* and Signal Recognition Particle are required for cardiac development and remodeling

**DOI:** 10.1101/2022.01.24.477419

**Authors:** Analyne M. Schroeder, Georg Vogler, Alexandre R. Colas, Rolf Bodmer

## Abstract

Congenital Heart Disease (CHD) is driven by a strong genetic predisposition, yet only a small subset of patients (∼20%) are diagnosed with a precise genetic cause. Therefore, expanding the pool of genes associated with CHD and establishing the functional relationships between genes can assemble a more comprehensive genetic network to better understand cardiac development and pathogenesis. In our studies, we identified protein biogenesis cofactors Nascent polypeptide Associated Complex (NAC) and Signal Recognition Particle (SRP) that bind disparate subsets of emerging nascent polypeptides at the ribosome exit site to direct polypeptide fates, as novel regulators of cell differentiation and cardiac morphogenesis. Knockdown (KD) of the alpha- *(Nacα)* or beta- subunit (*bicaudal, bic)* of NAC in the developing *Drosophila* heart led to disruption of cardiac remodeling during pupal stages resulting in an adult fly with no heart. Heart loss was rescued by combined KD of *Nacα* with the *Hox* gene *Abd-B.* Consistent with a central role for this interaction in the regulation of cardiogenesis, KD of *Nacα* in Cardiac Progenitors derived from human iPSCs impaired cardiac differentiation while co-KD with mammalian *Hox* genes *HOXC12 and HOXD12* rescued this phenotype. The effect of *Nacα* KD on the fly heart was temporally regulated, in that KD in embryo or in pupae caused only a partial loss of the heart, whereas KD during both stages led to heart loss similar to continuous KD throughout life. This suggests that *Nacα* KD already in the embryo may reprogram cells leading to aberrant cardiac remodeling during pupal stages. Lastly, KD of several SRP subunits individually in the fly heart produced a range of cardiac phenotypes that targeted specific segments and cell types, indicating spatially regulated activities of SRP components in the heart. Together, these data suggest that despite NAC and SRP ubiquitous presence, they displayed spatially and temporally fine-tuned activities for proper cardiac morphogenesis. *Nacα’s* interaction with cardiac-specific *Hox* gene functions builds upon the novel role of this pathway and expands our understanding of the complex genetic networks involved in cardiac development and pathogenesis.

## INTRODUCTION

Congenital Heart Disease (CHD) is characterized by structural malformations of the heart present at birth caused by deviations from the normal course of cardiogenesis [1]. Genetics is a critical driver of CHD [2, 3]. Chromosomal anomalies as well as variants in genes involved in heart development have been identified in CHD patients [4]. These genetic features and disease presentations are heritable and cluster in families [5, 6]. Identifying the genes associated with disease helps piece together genetic networks that could uncover mechanisms underlying pathogenesis. Approximately 400 genes have been implicated in CHD [2], some that cluster within defined pathways, which permits a genetic diagnosis for approximately 20% of CHD patients. However, this leaves the vast majority of CHD cases with unknown genetic origins [3]. Therefore, expansion of the genetic data pertinent to CHD, such as functional analysis of genes with variants of uncertain significance (VUS), would advance our understanding of the disease and may offer, in the future, a diagnosis and targeted treatment for CHD patients. A better understanding of additional genetic risk factors and patient-specific combinations of such factors is aided by identification of candidate disease genes through patient-specific genomics, such as whole genome sequencing (WGS), followed by their evaluation in cardiac developmental platforms and assays from various genetic model systems [7, 8]. These validation efforts can accelerate candidate gene identification and focus on new potentially pathogenic genes, including genes located within larger genomic anomalies such as *de novo* Copy Number Variants [9].

Previous data suggest that the human gene Nascent polypeptide Associated Complex-alpha (*NACA*) is a candidate CHD gene that could provide novel insights into biological pathways in cardiac morphogenesis and pathogenesis. Using a GWAS approach, *NACA* was located within a genomic locus associated with increased myocardial mass [10], while Whole Exome Sequencing in families with Tetralogy of Fallot identified a single nucleotide polymorphism within *NACA* [11]. *NACA* is a highly conserved *alpha* subunit of a heterodimeric complex called Nascent polypeptide Associated Complex (NAC). Along with its heterodimeric partner, *NAC*-*beta (NACβ/BTF3), NAC* is one of several chaperones found near the ribosome exit tunnel that bind to select emerging nascent polypeptides [12]. NAC-ribosome complexes facilitate transport of nascent polypeptides to the mitochondria as has been demonstrated in yeast [13–15]. At the ribosome exit site, NAC gates the activity of other nascent polypeptide chaperones. For example, NAC enhances the fidelity of Signal Recognition Particle (SRP) binding to only those nascent polypeptides destined for import to the Endoplasmic Reticulum (ER) [16–18]. Depletion of NAC leads to promiscuous binding of SRP onto nascent polypeptides, drawing mistargeted ribosome-nascent polypeptide complexes to the ER for aberrant insertion into the membrane or secretion. *NACA’s* function as part of NAC therefore regulates the localization and posttranslational quality control of proteins in the cell.

In *Drosophila,* NAC plays an important role in translational regulation critical for embryonic development [19, 20]. Fly homologs of NAC subunits, *Nacα* and *bicaudal (bic),* were shown to repress protein translation of the posterior patterning gene *Oskar* (*Osk)* in anterior regions of the embryo, which was despite association of *osk* mRNA with polysomes, usually indicative of active translation. Restricting Osk protein translation and accumulation to the posterior pole of the embryo is critically required for patterning the posterior body plan [21]. Depletion of either *Nacα* or *bic,* expands Osk protein localization anteriorly resulting in a bicaudal phenotype, where the embryo develops with mirror-image duplication of the posterior axis [19, 22]. These results suggest that within the developing embryo syncytium, NAC appears to regulate the expression of select proteins for proper spatial distribution. NAC could have a similar role in the timing and spatial targeting of translation within specific tissues.

*Nacα* has been demonstrated to be critical for development of several tissues using various model organisms. In mouse, the *Nacα* subunit can function as a transcriptional coactivator regulating bone development [23–25] and hematopoiesis in zebrafish [26]. In vertebrates, a skeletal muscle- and heart-specific variant of *Nacα* has been associated with myofibril organization in zebrafish [27] and muscle and bone differentiation in mouse [28–32]. Recent studies in the fly showed that *Nacα* knockdown (KD) specifically in the heart led to a ‘no adult heart’ phenotype [10], suggesting that *Nacα* could play a role in sarcomeric biogenesis, but its exact role in cardiac development had been unclear.

Here, we provide evidence for a cardiac developmental role for *NAC and SRP* in *Drosophila* and *NACA* in human Multipotent Cardiac Progenitors (MCPs). In flies, cardiac KD of *NACα* (mentioned above) and *bic* throughout development led to complete loss of the heart. We demonstrate that this phenotype is dependent on the timing of *NACα* KD, which requires KD during both embryonic heart development and pupal cardiac remodeling from larval to adult heart for a complete loss of the heart. KD in embryos only led to loss of the terminal chamber of the adult heart, while retaining heart structures in the anterior segments. KD of *Nacα* only during pupal stages did not affect adult heart structure. This suggests that NAC*α* KD primes cardiac cells already in the embryo for aberrant responses to morphogenic cues during later developmental stages. Persistent *Nacα* KD during pupal stages remained required for complete loss of the adult heart. Consistent with this idea, *NACα* KD throughout cardiac development induced ectopic expression of the posterior patterning *hox* gene *Abdominal-B* (*Abd-B*) into anterior regions of the remodeling heart during pupation. Concurrent KD of *NACα* and *Abd-B* partially rescued the heart. This interaction between *Nacα* and *Hox* gene was recapitulated in MCPs, whereby *NACA* KD led to deviations in progenitor cell differentiation away from cardiomyocytes and toward fibroblast cell fates, which was reversed with combined KD of *NACA* and *Hox* genes *HOXC12* an *HOXD12*. Because NAC associates and influences the activity of SRP, we tested the effects of individual SRP subunit KD on the fly heart. Interestingly, KD of individual SRP subunits produced cardiac phenotypes that were distinctly different from those of *NACα*, affecting other heart regions and targeting specific cell types. These results suggest specific roles for ubiquitously expressed protein biogenesis factors NAC and SRP in heart morphogenesis, in part through alterations in the expression of body patterning proteins. Translational regulation adds to our growing knowledge of biological pathways that may specifically contribute to cardiac pathogenesis leading to CHD.

## RESULTS

### Knockdown of *Nacα* and *bicaudal* in the heart throughout development results in complete absence of the adult heart

The Nascent Polypeptide Complex (NAC) is a heterodimeric complex made of an alpha (NACA/*Nacα)* and beta (NAC*β*/BTF3/bicaudal) subunit that bind ribosomes to influence translation, protein folding, and transport of select nascent polypeptide chains. We investigated the role of the NAC complex, focusing on the alpha subunit, in the development and function of the *Drosophila* heart. Consistent with previously published data [10], knockdown (KD) of *Nacα* by RNAi using the Hand4.2-GAL4 driver [33] that is active in the heart tube and surrounding pericardial cells throughout development, led to a ‘no-heart’ phenotype in adults when stained with phalloidin and the heart specific collagen, *pericardin* (**Figure 1A,B**). Similarly, KD of the *β*-subunit, *bicaudal*, using Hand4.2-GAL4 also led to a ‘no-heart’ phenotype (**Figure 1C**). Despite *Nacα*, or *bic* KD throughout heart development and the absence of hearts in adults, the posterior aorta and heart tube was still present in controls, *Nacα* KD, and *bic* KD during early pupae as captured by *in vivo* imaging of fluorescently labeled hearts (**Figure 1D-F**). We filmed and followed cardiac remodeling through pupation (**Supplemental Video 1-3, Supplemental Figure 1A**). Within the first 24 hours of pupation, hearts were detected in both controls and *Nacα* KD flies. Normally, the heart undergoes remodeling through trans-differentiation of the larval aorta into the adult heart tube (**Figure 3A**). However, after 24-28 hours of puparium formation (APF) the heart of cardiac *Nacα* KD flies began to disappear as cardiac remodeling progressed (**Supplemental Figure 1A**). The space at the dorsal midline usually taken up by the heart, is now filled in with fat cells, indicating that the heart undergoes complete histolysis during remodeling. A similar course of events is observed when *bicaudal* is knocked down in the heart (**Supplemental Video 3**; **Supplemental Figure 1A**). We also stained *Nacα* KD hearts with phalloidin just prior to cardiac remodeling (about 26-28 AFP), where heart tubes were present, but were narrower with disorganized actin filament arrangement (**Supplemental Figure 1C**).

**FIGURE 1:**
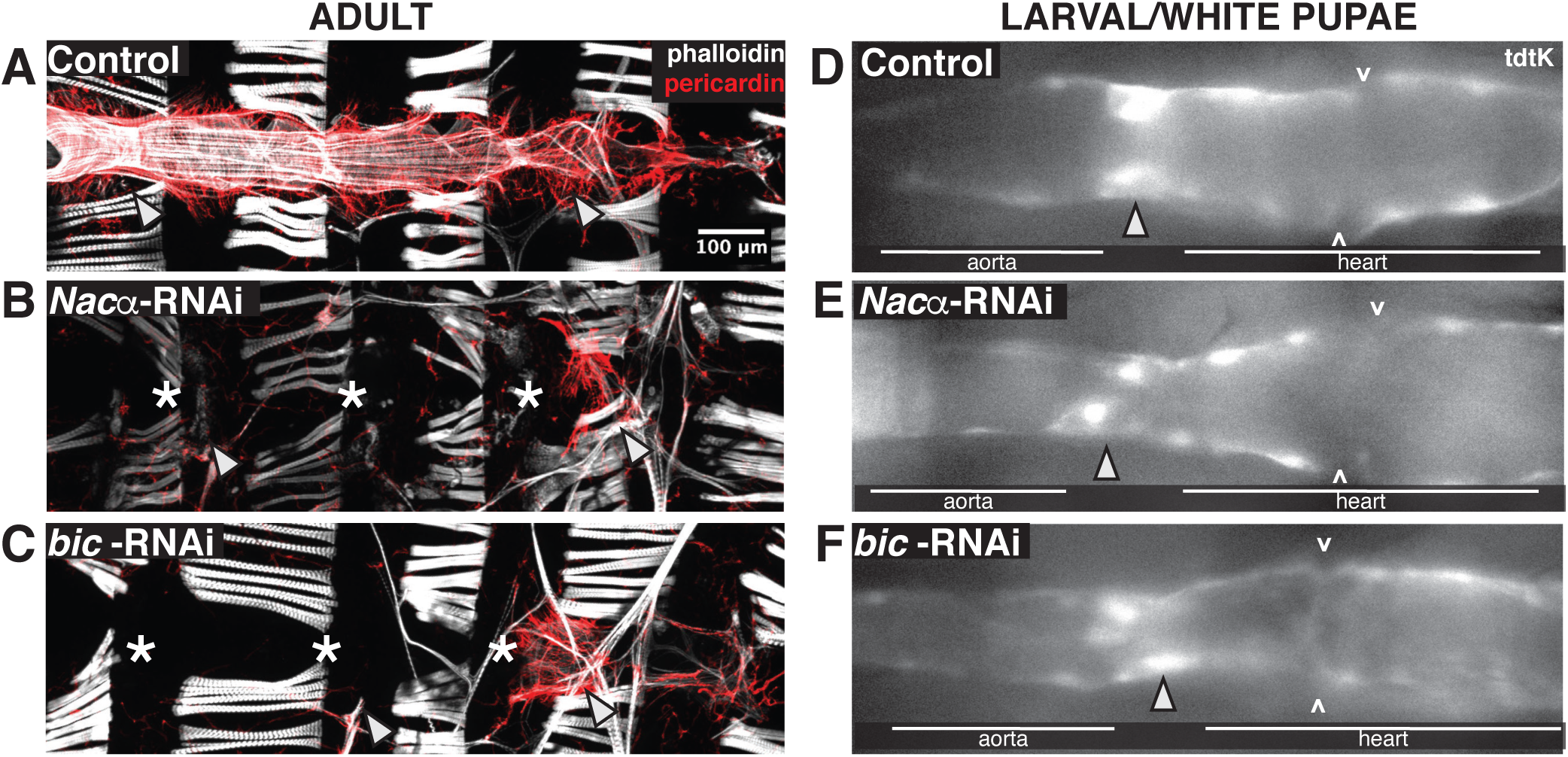
Knockdown (KD) of either subunit of Nascent polypeptide Associated Complex led to complete histolysis of the heart in adult flies. *Hand4.2*-GAL4 driving heart specific expression of **A,** control **B,** *Nacα-*RNAi or **C,** *bicaudal*-RNAi. KD of *Nacα or bicaudal* led to an absent heart in adult flies. * indicates absence of the heart tube. Arrowheads point to remnants of alary muscles that normally attach to the heart for structural support (anterior-left). Pericardin, a heart-specific collagen is largely absent, except for remnants in the posterior end. **D-F,** Fluorescently tagged hearts using tdtK were imaged *in vivo* at white pupae stages which displayed structurally intact hearts in **D,** control, **E,** *Nacα-*RNAi and **F,** *bicaudal*-RNAi expressing hearts, suggesting that the heart histolyzes during cardiac remodeling during metamorphosis. Arrow heads point to the internal valves that separate the larval aorta from the heart. ^ point to the inflow tracts called ostia.

Using cardiomyocyte-specific drivers conferring weaker expression than Hand4.2-GAL4 to reduce *Nacα* levels, *i.e. tinHE*-GAL4 and *tinCΔ*4-GAL4, did not produce severe heart loss but altered several parameters of heart function and structure, as measured by SOHA (see methods; **Figure 2A,B**). Diastolic diameters were decreased using both drivers (**Figure 2C**) without a change in systolic diameter (**Figure 2D**), and consequently reducing contractility, as measured by diminished fractional shortening, FS (**Figure 2E**). No changes were detected in heart period (**Figure 2F**) and diastolic intervals (**Figure 2G**). KD of *Nacα* with *tinCΔ*4-GAL4 caused a slight reduction in systolic interval, which is the duration of active heart contraction and relaxation (**Figure 2H**). Phalloidin staining of the adult hearts with reduced *Nacα* expression showed gaps between the circumferential myofibrils and sarcomeric disorganization compared to controls, consistent with reduced contractility (**Figure 2I**). These results suggest that at certain expression thresholds where heart development is not completely diverted, *Nacα* may play a role in functional and structural aspects of the heart as well. Because *Hand*4.2-GAL4 drives expression in both cardiac cells and the pericardial nephrocytes, we tested the effect of *Nacα* KD in pericardial cells only using *Dot*-GAL4 [34] to assess their contribution to the overall heart phenotype (**Figure 2 B-H**). KD in the pericardial cells did not produce significant changes in heart function, except for a slight increase in systolic diameter but no change in contractility (**Figure 2D,E**). The organization of the circumferential myofibrils as stained by phalloidin was unaltered compared to control (**Figure 2I**), suggesting that KD of *Nacα* in pericardial cells contribute minimally to the overall cardiac phenotype.

**FIGURE 2:**
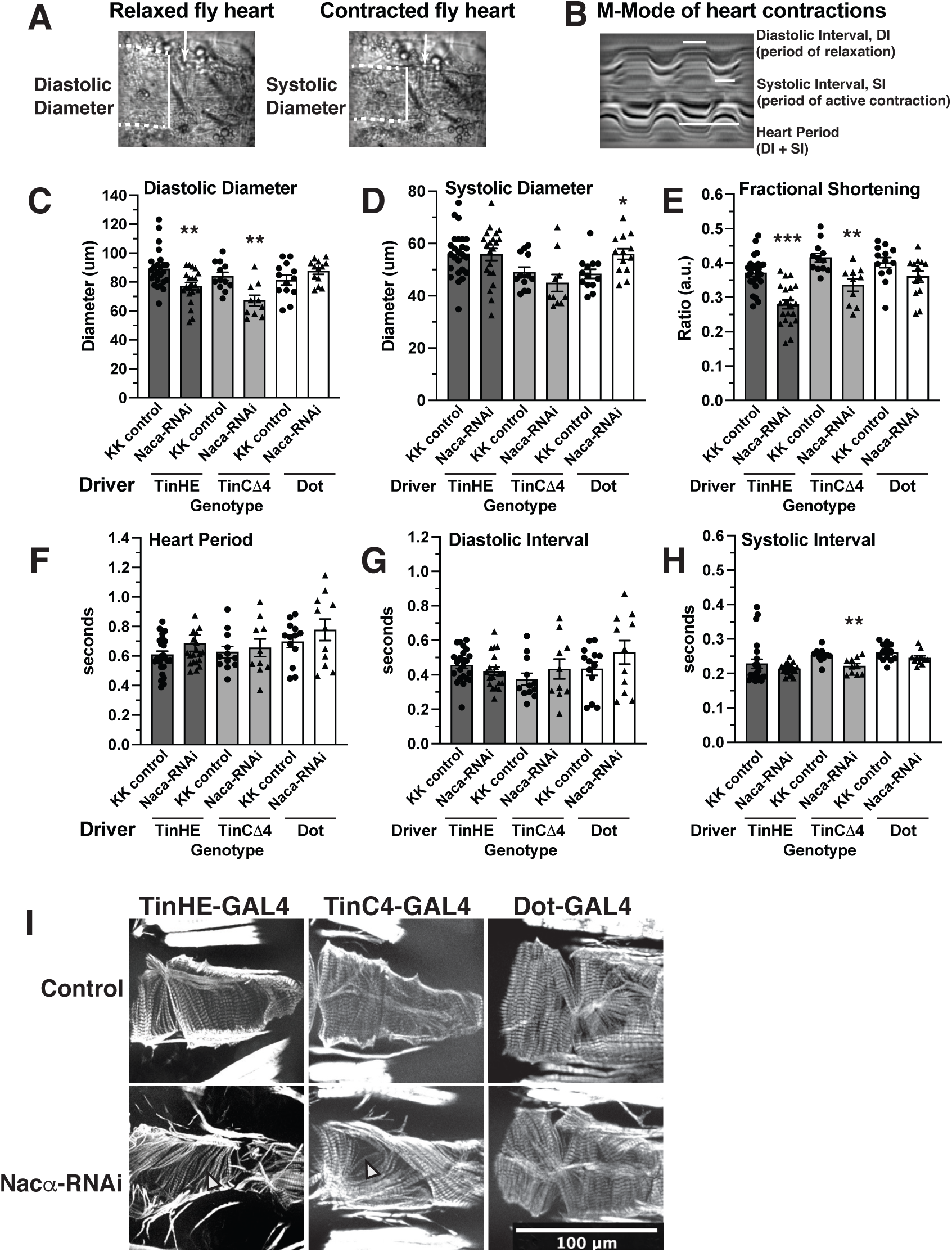
Effect of *Nacα* knockdown (KD) using various GAL4 drivers on heart function and structure. **A**,**B** Structural and functional parameters measured by SOHA to assess the fly heart. Dotted lines indicate the heart tube borders. White solid vertical lines indicate the diameters of the heart, while horizontal lines indicate the duration of contraction/relaxation being measured. **A,** Diastolic Diameter measures the heart diameter when it is fully relaxed, while systolic diameter measures the heart diameter when it is fully contracted. **B**, Motion-mode (m-mode) of the heart for temporal resolution of heart movement. Diastolic Interval measures the duration during which the heart is non-contractile, which occurs in this denervated fly preparation. Systolic Interval measures the duration that the heart is in active contraction and relaxation. **C**, Both *tinHE*-GAL4 and *tinCΔ4*-GAL4 cardiac drivers reduced Diastolic Diameter when used to KD *Nacα* expression. *Dot*-GAL4 pericardial cell driver had no effect on diastolic diameter. **D,** Cardiac drivers had no significant effect on systolic diameters, while *Dot*-GAL4 increased systolic diameter slightly, indicating mild systolic dysfunction. **E,** Fractional Shortening is significantly decreased using both *tinHE*-GAL4 and *tinCΔ4*-GAL4 driver, while *Dot*-GAL4 had no effect. No changes in Heart Period **F,** or Diastolic Interval **G,** were detected with either cardiac drivers or *Dot*-GAL4. **H,** A slight reduction in systolic interval was detected when *Nacα*-RNAi was driven with *tinCΔ4*-GAL4. **I,** Adult fly hearts were stained with phalloidin to examine cytoskeletal structures following *Nacα* KD using various tissue drivers. Compared to controls that display well- and tightly organized circumferential fibers that drive heart contractions, both *tinHE*-GAL4 and *tinCΔ4*-GAL4 drivers led to alterations in the organization of fibers. White arrowheads point to gaps in the fibers in KD samples consistent with the observed reductions in fractional shortening. KD of *Nacα* expression using *Dot*-GAL4 did not cause alterations in circumferential organization.

### Nac*α* genetically interacts with the *Hox* gene *Abd-B* in heart development

We sought to better understand the mechanisms driving complete histolysis of the heart tube during metamorphosis. It is well established that during normal cardiac remodeling the posterior most segment of the larval heart (abdominal segments 6-7) histolyzes and is no longer present in adult hearts [35]. The posterior larval heart region destined for histolysis expresses the *Hox* segmentation gene *Abdominal-B* (*Abd-B*) [36–38]. Abdominal segment 5 in the larvae that expresses the *hox* segmentation gene *abdominal-A* (*abd-A*) remodels to become the adult terminal chamber, while the larval aorta (abdominal segment 1-4) that expresses Hox gene *Ultrabithorax (Ubx)* remodels to become the adult heart proper (**Figure 3A**). We postulated that KD of *Nacα* throughout the heart could result in the misexpression of *Abd-B* leading to complete histolysis of the heart. First, we examined whether overexpression of *Abd-B* throughout the heart would result in similar phenotypes as with *Nacα* KD. Previously published data overexpressing *Abd-B* using an early pan-mesodermal driver (*twist*-GAL4) led to severely diminished embryonic muscle and heart development [39]. Using the cardiac-specific driver Hand4.2-GAL4 to overexpress *Abd-B*, the heart was completely absent in adults (**Figure 3B, right**), however a heart tube was present in larvae and in early pupae (24 hours APF, **Figure 3D**), similar to control (**Figure 3C**) *Nacα-RNAi* (**Figure 3E**) and *bic-RNAi* phenotypes (**Figure 1**). These results suggest that *Abd-B* expression and activity are highly temporally controlled, and drive cardiomyocyte histolysis only during cardiac remodeling at pupal stages.

**FIGURE 3:**
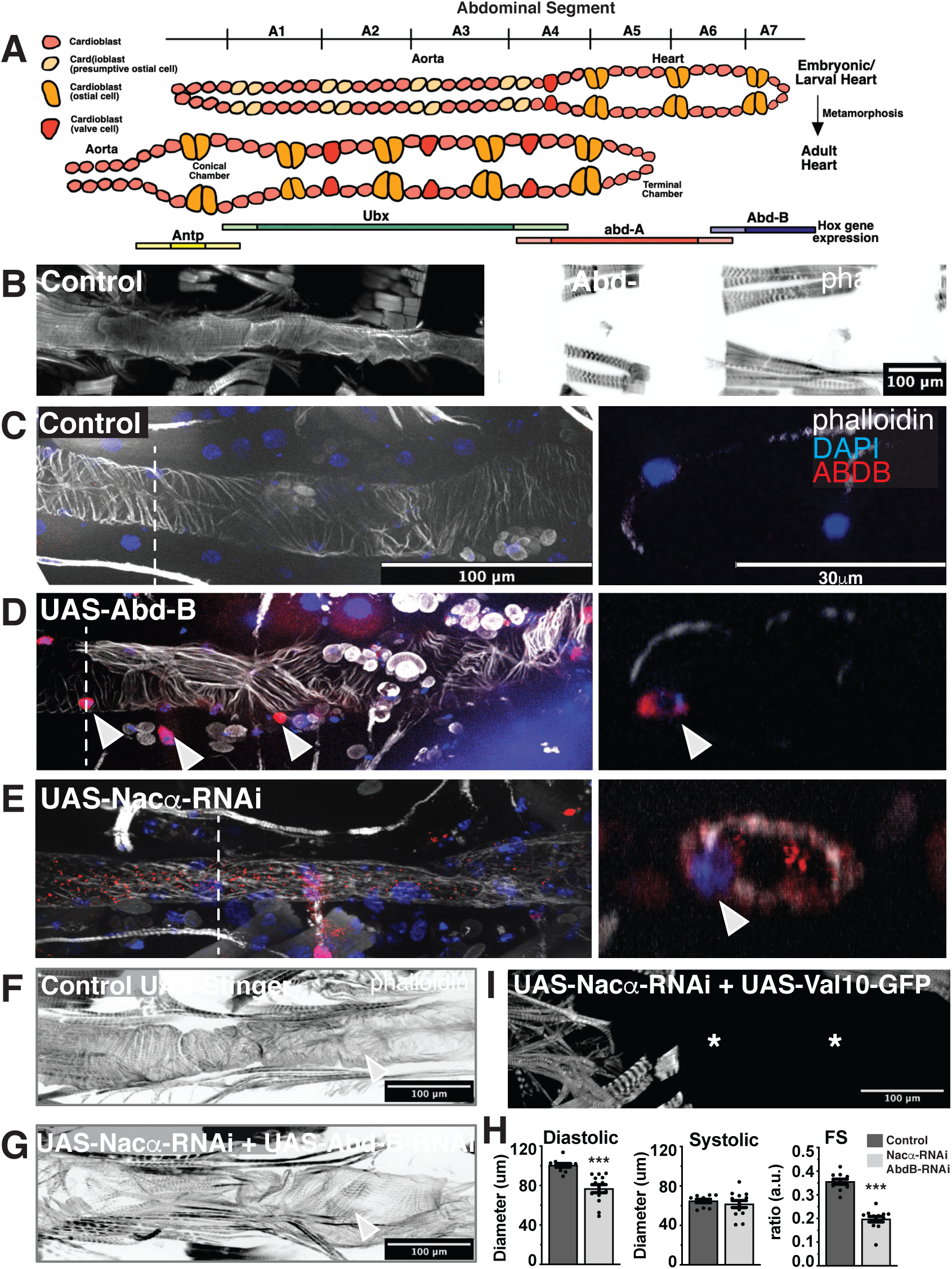
*Nacα* and the *Hox* gene *Abdominal B* (*Abd-B*) genetically interact in the heart. **A,** The embryonic/larval fly heart remodels into adult structures through trans-differentiation of the *Ubx* and *abd-A* expressing cardiomyocytes into the adult heart and terminal chamber, respectively. The cardiomyocytes of the posterior larval heart that express *Abd-B* (abdominal segment 6-7) histolyze and are absent in the adult heart. **B,** Overexpression of *Abd-B* in the heart using the cardiac driver *Hand4.2*-GAL4 led to complete absence of the adult heart, which in early stages of pupation was still present (see **D**). **\*** indicates absence of the heart tube. **C-E.** Immunohistochemistry of pupal hearts 26-28 hours After Puparium Formation (APF). Dashed lines mark the region of the heart where orthogonal cross-sections were taken to examine Abd-B and DAPI expression in the nuclei. Images of the cross-section of cardiomyocytes and nuclei are displayed in the right panel. Arrowheads indicate either cardiac and pericardial nuclei that are both Abd-B and DAPI positive. **C,** In controls, phalloidin stained the circumferential fibers of the pupal aorta and heart. Abd-B staining was not detected in the cardiomyocyte nuclei within aorta and anterior heart segments but, ABDB is stained in the posterior segments of the embryo. **D,** Overexpression of *Abd-B* using *Hand4.2*-GAL4 resulted in strong Abd-B staining in the nuclei (as indicated by the arrowheads) throughout the pupal heart and pericardial cells prior to remodeling. Cross section clearly shows that Abd-B is localized in the nucleus. **E,** Knockdown of *Nacα* in the heart resulted in ectopic Abd-B expression throughout the heart tube. Staining was present within the nuclei (marked by DAPI), cytoplasm and heart lumen (see orthogonal sections, right). **F-H.** Concurrent knockdown of *Nacα and Abd-B* in the heart, **G**, led to rescue of heart tube formation, with visible circumferential fibers (arrowheads) albeit less well-organized than controls. Rescued hearts contracted as visualized in the M-mode (right panel). **H,** Rescued hearts were smaller in diastolic diameter with no change in systolic diameter resulting in reduced Fractional Shortening (FS). This effect was not due to titration of GAL4 onto 2 UAS sites, as combination of UAS-*Nacα-RNAi* with UAS-Val10-GFP still produced a no heart phenotype. **I**, Arrow indicates remnants of the ventral longitudinal muscle in the anterior end of the abdomen.

We then examined whether KD of *Nacα* leads to ectopic expression of Abd-B protein in the early pupal heart just prior to remodeling (APF 26-28). In controls, Abd-B protein is undetected in the anterior regions of the early pupal heart tube (**Figure 3C**), but detected in the posterior segments of the embryo. Overexpression of *Abd-B* using *Hand*4.2-GAL4 led to increased Abd-B protein accumulation throughout the early pupal heart tube and pericardial cells, which was restricted to the nuclei (**Figure 3D**). When *Nacα* was knocked down in the heart, we detected Abd-B protein expression throughout the larval/early pupal heart tube, within the myocardial nuclei as well as in the cytoplasm (**Figure 3E**). In orthogonal sections, there is ectopic Abd-B staining centrally, possibly within the lumen of the heart, either secreted from cardiomyocytes or bound to circulating hemocytes. These data suggest that KD of *Nacα* in the heart led to ectopic expression of Abd-B within the cell, as well as, anteriorly throughout the heart tube. Because overexpression of Abd-B using *Hand*4.2-GAL4 led to heart histolysis during remodeling, we speculate that ectopic Adb-B expression induced by *Nacα* KD is likely leading to the observed histolysis of the entire heart during remodeling.

Because *Nacα* KD led to ectopic expression of Abd-B, we tested whether the no-heart phenotype produced by *Nacα* KD could be rescued by concurrent KD of *Abd-B*. KD of *Abd-B* alone led to largely intact hearts except for posterior ends that were more dilated and prominent compared to controls suggesting incomplete histolysis during cardiac remodeling **(Supplemental Figure 2**). KD of both *Nacα* and *Abd-B* using the Hand4.2-GAL4 driver led to a remarkable rescue of the adult heart, restoring formation of most of the circumferential myofibrils in the anterior regions (**Figure 3F,G**). These hearts had reduced diastolic diameters, without change in systolic diameter that significantly reduced FS (**Figure 3H**). This rescue was not due to titration of GAL4 protein by competing UAS sites, since a no-heart phenotype was still produced when *Nacα-RNAi* was combined with another UAS that carries a UAS-Val10-GFP construct (**Figure 3I**). These rescue data suggest a genetic interaction between *Nacα* and *Abd-B*. On the other hand, co-expression of *Nacα-RNAi* with an inhibitor of apoptosis (Death-associated inhibitor of apoptosis 1, DIAP1) did not rescue the heart (**Supplemental Figure 3**), suggesting that the *Nacα*-RNAi mediated heart loss cannot simply be prevented by inhibition of canonical cell-death pathways.

We also tested whether *Abd-B* KD could rescue the moderate cardiac function defects produced by less robust KD of *Nacα*, using the *tinHE*-GAL4 driver, that only drives expression during embryonic development. *Nacα* KD with *tinHE*-GAL4 caused cardiac constriction but no effects on heart period or intervals (**Supplemental Figure 4A,B)**. KD of *Abd-B* using *tinHE*-GAL4 did not produce significant changes in heart function compared to controls, except for a prolonged systolic interval. Combining *Nacα-RNAi* with *Abd-B-RNAi* reversed the constricted diastolic and systolic diameter phenotype of *Nacα-RNAi;UAS-Stinger::GFP*, such that the phenotype mirrored controls or *Abd-B-RNAi;UAS-Stinger KD*. An improvement in circumferential myofibrillar organization is also evident with co-KD of *Nacα and Abd-B* (**Supplemental Figure 4C**). These data suggest that *Abd-B and Nacα* co*-*KD cannot only restore heart formation with the strong *Hand4.2* driver, but also normal heart function and myofibrillar organization with the weaker *tinHE*-GAL4 driver. This means that functional maturation requires *Nacα* to restrict *Abd-B* function to the posterior of the heart.

### Nac*α* is required in the embryo to pre-program cardioblasts for appropriate cardiac remodeling

Prior to pupal cardiac remodeling, a larval heart was still present despite *Nacα* KD using the strong cardiac driver *Hand4.2*-GAL4 (**Figure 1E** **and 3E**). However, these larval heart tubes were thinner and the cytoskeletal structures less prominent than controls (**Supplemental Figure 1B and Figure 3E**), reminiscent to the constricted adult heart phenotypes with the weaker driver *tinHE*-Gal4 (**Supplemental Figure 4**). These observations suggest that *Nacα* may have earlier developmental functions in addition to a role in metamorphosis. We therefore wanted to temporally dissect *Nacα’s* function in the heart by knocking down its expression during different developmental stages. We generated a *Hand4.2*-GAL4 driver line that included two copies of a temperature-sensitive allele of GAL80 driven by a ubiquitous promoter (*tubulin*-GAL80^ts^), which we termed HTT [9]. At the permissive temperature (18°C), the GAL80 transcriptional repressor prevents GAL4 activation of UAS sites thereby inhibiting transcription of downstream constructs [40]. This temperature-sensitive form of GAL80 protein is unstable at higher temperatures (28-29°C), thus permitting GAL4 activity at higher ambient temperatures.

Maintaining HTT flies crossed to *Nacα-RNAi* or controls at 18°C throughout development resulted in normal heart structure (**Figure 4A**) and produced no differences in diastolic diameter, systolic diameter or fractional shortening compared to controls (**Figure 4I**), effectively demonstrating GAL80’s ability to suppress Nac*α*-RNAi transcription at 18°C temperatures. Constant exposure to 28°C throughout development phenocopies the absence of the heart in adults produced by Hand4.2-GAL4 driver (**Figure 4B**). KD of *Nacα* in adults only for one week by exposure to high temperatures also led to largely normal heart structure and function compared to controls (**Figure 4C,I**), suggesting that *Nacα* is primarily required developmentally for establishing a normal heart in adults, rather than maintaining its function or structure with age. Remarkably, although lifelong cardiac *Nacα* KD using Hand4.2-GAL4 led to histolysis of the heart during metamorphosis, KD of *Nacα* only during pupation did not produce gross heart defects (**Figure 4D**). The diastolic and systolic diameters were slightly increased, which caused some reduction in fractional shortening (**Figure 4I**). Even when we induced *Nacα* KD earlier, starting at mid-larval stages through metamorphosis until eclosion (∼5 days), a period with substantial developmental growth requiring high levels of protein translation, heart structure was unaffected (**Figure 4E**). This lack of phenotype is remarkable, as disruption of NAC is associated with proteostasis and ER stress, which often leads to cell death [41–43]. These results suggest that KD of *Nacα* in the developing heart, even for relatively long durations does not unequivocally lead to cell death. When *Nacα* was knocked down during embryonic stages only (egg-lay up to 24 hours) and subsequently reared at 18°C until dissection to prevent *Nacα* KD at later stages, most adult hearts remained intact, but considerably constricted with smaller diastolic and systolic diameters (**Figure 4F,I**). Interestingly, the posterior terminal heart chamber was absent in most cases (**Figure 4F**), which suggests that the no-heart phenotype observed with continuous KD likely arises from developmental defects already in the embryo. Extending exposure to higher temperature during embryonic stages until 48 hours after egg lay, produced similar phenotypes compared to 24hr exposure, with the presence of a heart tube but without a terminal chamber (**Figure 4G**). Remarkably, when flies were exposed to higher temperatures during both embryonic stages (24 hours) and pupal stages (three days), with a return to 18°C during larval development, an almost complete ‘no-heart’ phenotype was reproduced in most cases (**Figure 4H**). This suggests that *Nacα* KD only during combined embryonic and pupal stages produces a nearly complete loss of heart structures, but at either stage alone was insufficient for such a severe phenotype.

**FIGURE 4:**
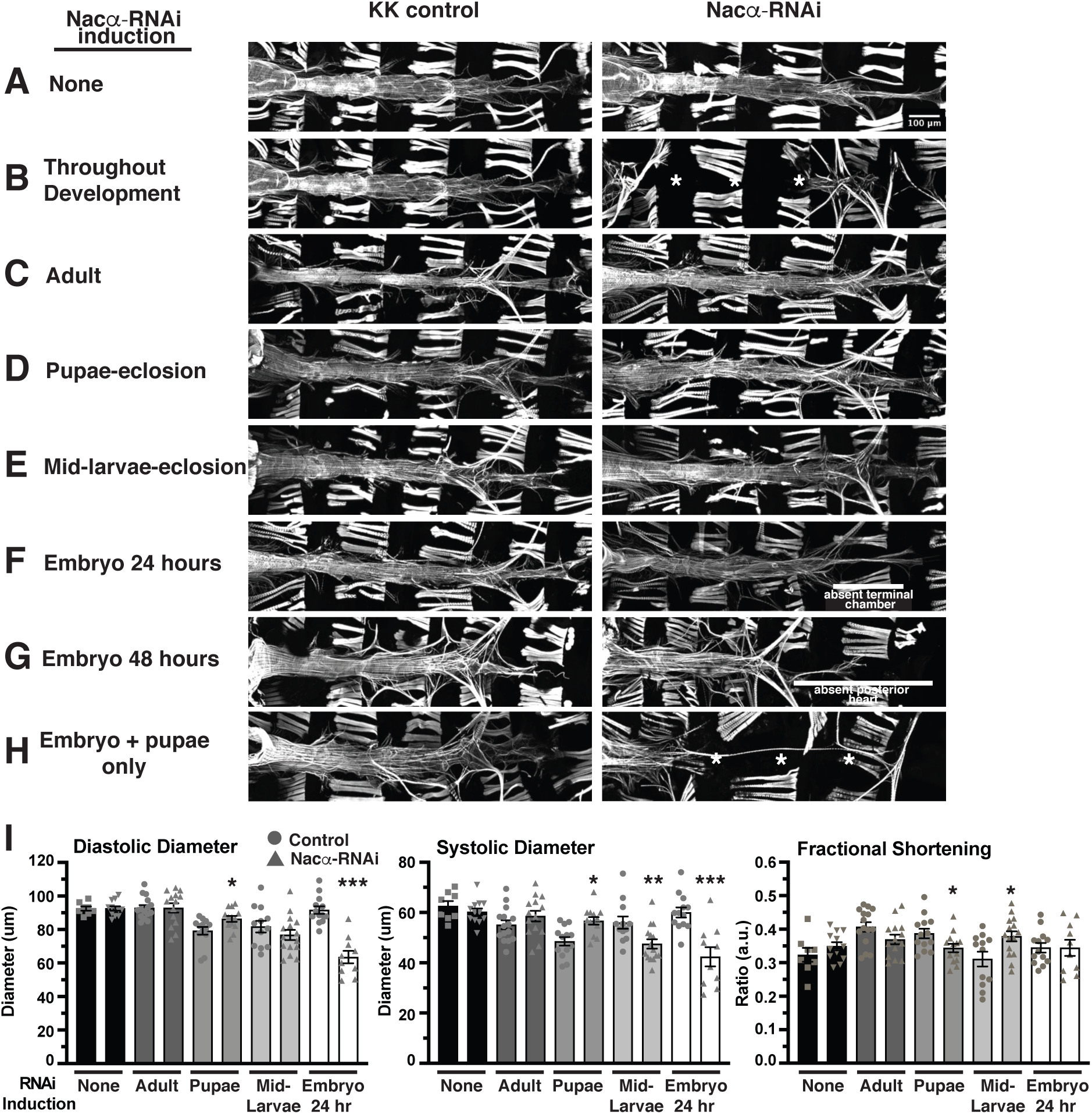
Temporal regulation of *Nacα-*RNAi expression in the heart. Using a temperature inducible driver specifically in the heart (HTT, *Hand4.2*-GAL4, *tubulin*-GAL80^ts^; *tubulin*-GAL80^ts^), *Nacα-*RNAi was expressed during specific stages of development by controlling ambient temperature to determine its contribution to cardiogenesis. Controls lacking RNAi are kept in similar temperature conditions to account for any developmental effects of temperature on the heart. **A-H.** Phalloidin staining to visualize cytoskeletal structural effects of *Nacα* knockdown (KD). **A,** As a test of GAL80 control of transcription, flies held at 18°C throughout development did not produce changes to heart structure indicating an inhibition of *Nacα*-RNAi transcription. **B,** Exposing flies to high temperatures (28°C) throughout development produced a no heart phenotype similar to the effects of driving *Nacα-RNAi* using *Hand4.2*-GAL4 alone, suggesting an induction of *Nacα*-RNAi transcription and subsequent *Nacα* KD with exposure to higher temperatures. * indicates absence of the heart. **C,** Exposing flies to high temperature during adulthood only for 1 week, **D,** pupae to eclosion, or **E,** mid-larvae to eclosion did not produce gross structural defects in the heart. **F,** Exposing embryos to high temperatures starting at egg-lay up until 24 hours resulted in the absence of the terminal chamber, indicated by white bar. Only thin alary muscles were present. **G,** Extending the high temperature exposure to 48 hours led to similar loss of the posterior heart, indicated by white bar. **H,** Only when the hearts were exposed to higher temperatures during embryonic stage (24 hours) and pupal stage until eclosion, were we able to recapitulate the no heart phenotype produced by exposing the heart to constant high temperatures. **I,** Functional analysis of the adult heart following *Nacα* KD at various developmental stages. Maintaining flies at 18°C throughout development or exposure of adult flies to high temperature for 1 week led to no changes in diastolic diameter, systolic diameter, or fractional shortening. High temperature exposure from pupae to eclosion or from mid-larvae to eclosion led to subtle changes in diameters and fractional shortening. Exposure of embryos to high temperatures for 24hr led to constricted diastolic and systolic diameters. * *p<0.05, ** p<0.01, *** p<0.001*.

These results suggest that *Nacα’s* role in driving heart morphogenesis has a temporal component, perhaps regulating several different processes during cardiac development. Furthermore, the observation that *Nacα* KD during embryonic as well as pupal stages is needed to induce histolysis of nearly the entire heart tube during pupal cardiac remodeling suggests that *Nacα* plays an essential role in the embryonic heart development by programing cardiac cell fate. This embryonic requirement seems to be partially compensated for by *Nacα* function during heart remodeling. However, the additional reduction of pupal *Nacα* exacerbates the lack of embryonic cardioblast programming, together leading to a failure of the larval heart to respond to remodeling cues during metamorphosis, thus causing histolysis.

### Nac*α* alters cell-fate in human Multipotent Cardiac Progenitors and is modulated by *Hox* genes

Since our results suggest that *Nacα* plays a role in establishing cell identity and cell-fate of cardiac cells in *Drosophila*, we wanted to determine whether a similar role could be observed in other model systems, such as cardiomyocytes derived from human iPSCs (hiPSCs). We therefore subjected human iPSC-derived Multipotent Cardiac Progenitors (MCPs)[44] to siRNAs against the human ortholog of *Nacα*, *NACA,* to assess their effects on cell proliferation and differentiation (**Figure 5**)[9]. We evaluated the propensity for MCPs to spontaneously differentiate into different cell types and calculated their relative proportions, by staining with *α-Actinin1* (*ACTN1*) for cardiomyocytes (CM), *Transgelin* (*TAGLN*) for fibroblasts, and *Cadherin 5* (*CDH5*) for endothelial cells. Total cell number was quantified by counting the number of DAPI-positive nuclei. The proportion of differentiated cell types was assessed nine days after siRNA treatment, a timepoint when active KD is no longer expected [9]. Efficiency of siRNA KD following transfection is expected to be maintained even when siRNAs against 3 different gene targets are combined, as demonstrated in previous experiments [7, 9].

**FIGURE 5:**
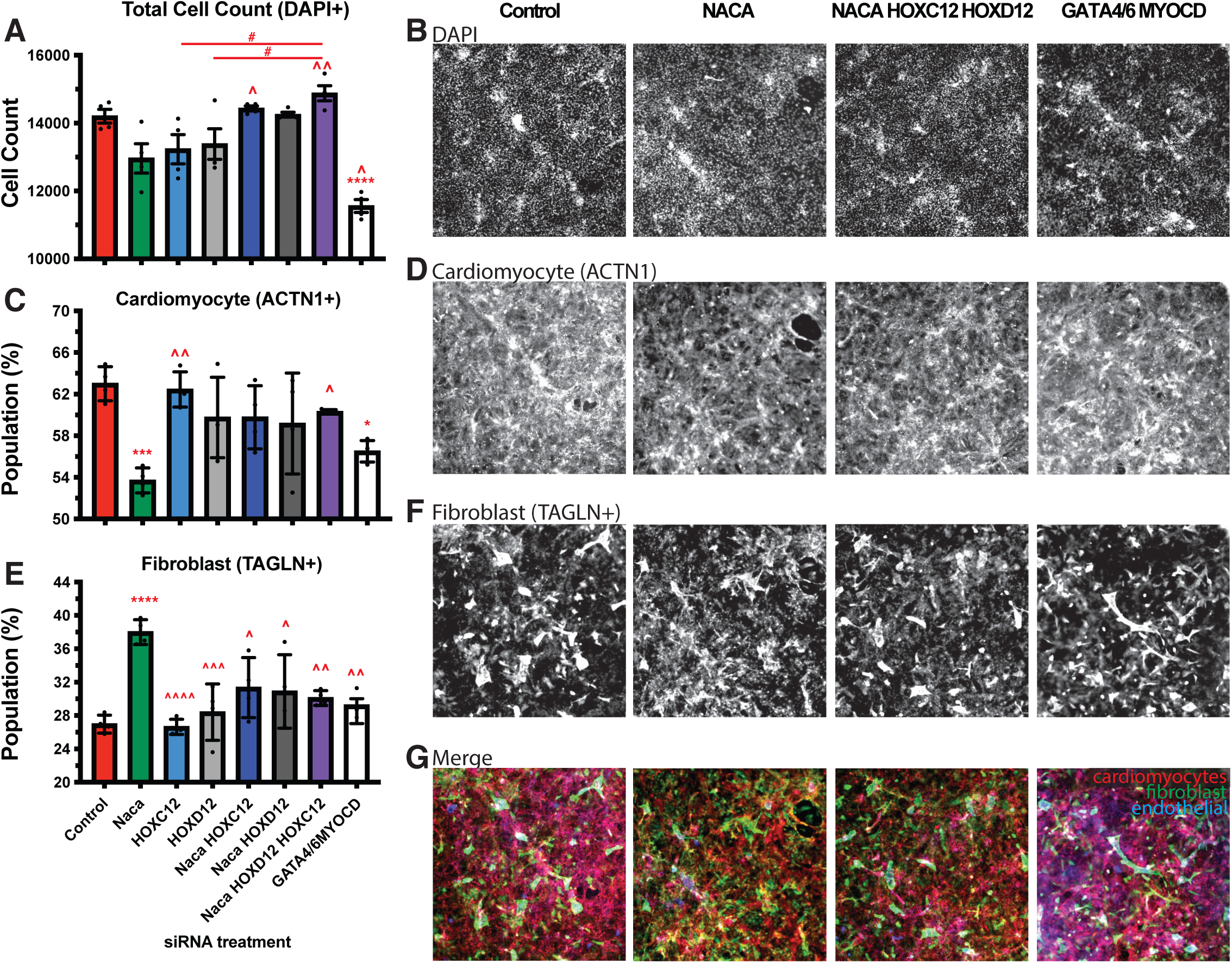
*Nacα* and *Hox* genes interact to redirect differentiation of Multipotent Cardiac Progenitors (MCPs). **A,C,E,** Quantitation of differentiated cell populations 9 days after siRNA treatment. **B,D,F.** Representative images of immunohistological staining for select conditions. **A,B,** Total Cell Populations following siRNA treatment were not significantly changed compared to controls, except for *Gata4/6,MyoCD* siRNA condition which reduced overall cell count. Knockdown (KD) of *Nacα* (green), *HOXC12* (light blue)*, HOXD12* (light gray) singly resulted in cell populations that trended lower. This decrease was reversed and significantly different upon combined transfection of *Nacα siRNAs* with *Hox* genes, compared to single siRNA transfections. **C,D,** *Nacα* KD alone significantly decreased the proportion of cardiomyocytes (ACTN1+) compared to controls, while treatment with HOXC12 or HOXD12 siRNA individually, had no effect. Combined transfection of *Nacα* and *Hox* genes reversed the decrease in cardiomyocyte population and was significantly different compared to *Nacα* KD alone and no longer different compared to controls. *Gata4/6,MyoCD* KD also significantly lowered the proportion of cardiomyocyte populations. **E,F,** *Nacα* KD increased the proportion of fibroblasts (TAGLN+) compared to controls. Combined KD of *Nacα* with any of the *Hox* genes did not alter fibroblasts numbers compared to controls but were significantly reduced compared to *Nacα* KD alone. **G,** Images of merged staining of cardiomyocyte, fibroblast, and endothelial cells. Significance * vs. control. ^ vs. *Nacα. #* comparison is indicated by line. * *p<0.05, ** p<0.01, *** p<0.001,* **** *p<0.0001*.

Treatment of MCPs with *NACA siRNA* did not change total cell count compared to controls (**Figure 5A,B**), but significantly decreased the proportion of CMs (**Figure 5C,D**), and increased the proportion of fibroblast (**Figure 5E,F**). The proportion of endothelial cells remained unchanged **(Supplemental Figure 5A,B)**. These results suggest that *NACA* may play a role in directing cell fate toward a cardiac program and its absence shifts these fates toward fibroblast differentiation.

We wondered whether the effect of *NACA* on CM differentiation can be similarly reversed by human orthologs of *Drosophila Abd-B*. Therefore, we tested the effect of siRNAs against human *Homeobox C12* (*HOXC12*) or *Homeobox D12* (*HOXD12*), which have close sequence homology with *Abd-B.* Treatment of either *HOXC12 and HOXD12 siRNAs* individually did not have a significant effect on total (**Figure 5A**), cardiomyocyte (**Figure 5C**), fibroblast (**Figure 5E**), or endothelial **(Supplemental Figure 5A)** cell populations compared to controls. When we tested combinations of *NACA* siRNA with *HOXC12* and *HOXD12* siRNA, or all three, we found that the *Hox* genes could alter *NACA* phenotypes. While treatment with *NACA* siRNA led to a reduced trend in total cell populations, combined KD of *NACA* and *HOXC12* and *HOXD12* led to total cell populations that were significant higher than *NACA siRNA* treatment alone and were more similar to controls (**Figure 5 A,B**). More striking is the reversal in the proportion of cardiomyocyte and fibroblast populations when *NACA siRNA* was combined with *HOXC12* and *HOXD12.* Co-transfection of *NACA* with the selected *Hox* genes led to increased CM (**Figure 5C,D**) and reduced fibroblasts (**Figure 5E,F**) such that they are significantly different compared to *NACA* siRNA treatment alone and no longer different compared to controls. These results suggest that the reduction in CM and increase in fibroblast caused by *NACA* KD is dramatically rescued upon *HOX* co-KD, thus restoring CM differentiation of these pluripotent cells (**Figure 5G**). These results are consistent with observed cardiac differentiation and genetic interactions in *Drosophila* and that *Nacα/NACA* activity in the heart may in part be mediated by posterior Hox genes.

As a positive control and a means of comparison, we transfected MCPs with siRNAs against cardiogenic transcription factors GATA Binding Protein 4 and 6 (*GATA4/6*) and Myocardin (*MYOCD*) [45, 46]. *GATA4/6,MYOCD* KD caused a significant decrease in total cell number (**Figure 5A,B**) and proportion of cardiomyocytes (**Figure 5C,D**), no change in the proportion of fibroblast (**Figure 5E,F**), and an increase in the proportion of endothelial cells (**Supplemental Figure 5A,B**). Thus, the effect of *NACA* KD on the proportion of cell types were different compared to the cardiogenic factors. The only similarity observed between *NACA* and *GATA4/6,MYOCD* KD was a decrease in the proportion of cardiomyocytes, although *NACA* produced a greater decrease (**Figure 5 C,D**). These results suggest that *NACA KD* induced alternate cell fates, possibly by a different mechanism to that of these cardiogenic transcription factors.

### Knockdown of individual SRP subunits cause distinct cardiac phenotypes

NAC is just one of the protein biogenesis quality control mechanisms that is found at the ribosomal exit site sifting through emerging nascent polypeptides and guiding protein fates. SRP is another protein complex involved in controlling protein biogenesis by binding a disparate subset of nascent polypeptides destined for the ER, wherein its targeting is influenced by the activity of NAC. Therefore, we wanted to examine if disruption of SRP function in fly hearts would result in similar defects compared to *Nacα* KD. Eukaryotic SRP is composed of a 7SL SRP RNA that holds the conformation of 6 proteins (SRP9, SRP14, SRP19, SRP54, SRP68, and SRP72) and targets approximately 30% of newly synthesized proteins to the ER (**Figure 6A**)[47]. SRP primarily recognizes the N-terminal hydrophobic sequences of emerging nascent polypeptides, but has also been shown to bind nascent chains even when target sequences are not yet accessible [48]. In yeast, SRP also binds nascent chains with internal transmembrane domains [49]. Once bound, SRP arrests translation of the nascent chain until the SRP-ribosome complex binds with the SRP-receptor (SR) anchored to the ER, where translation of the nascent chain is restarted and co-translationally released through the Sec61p translocase for insertion.

**FIGURE 6:**
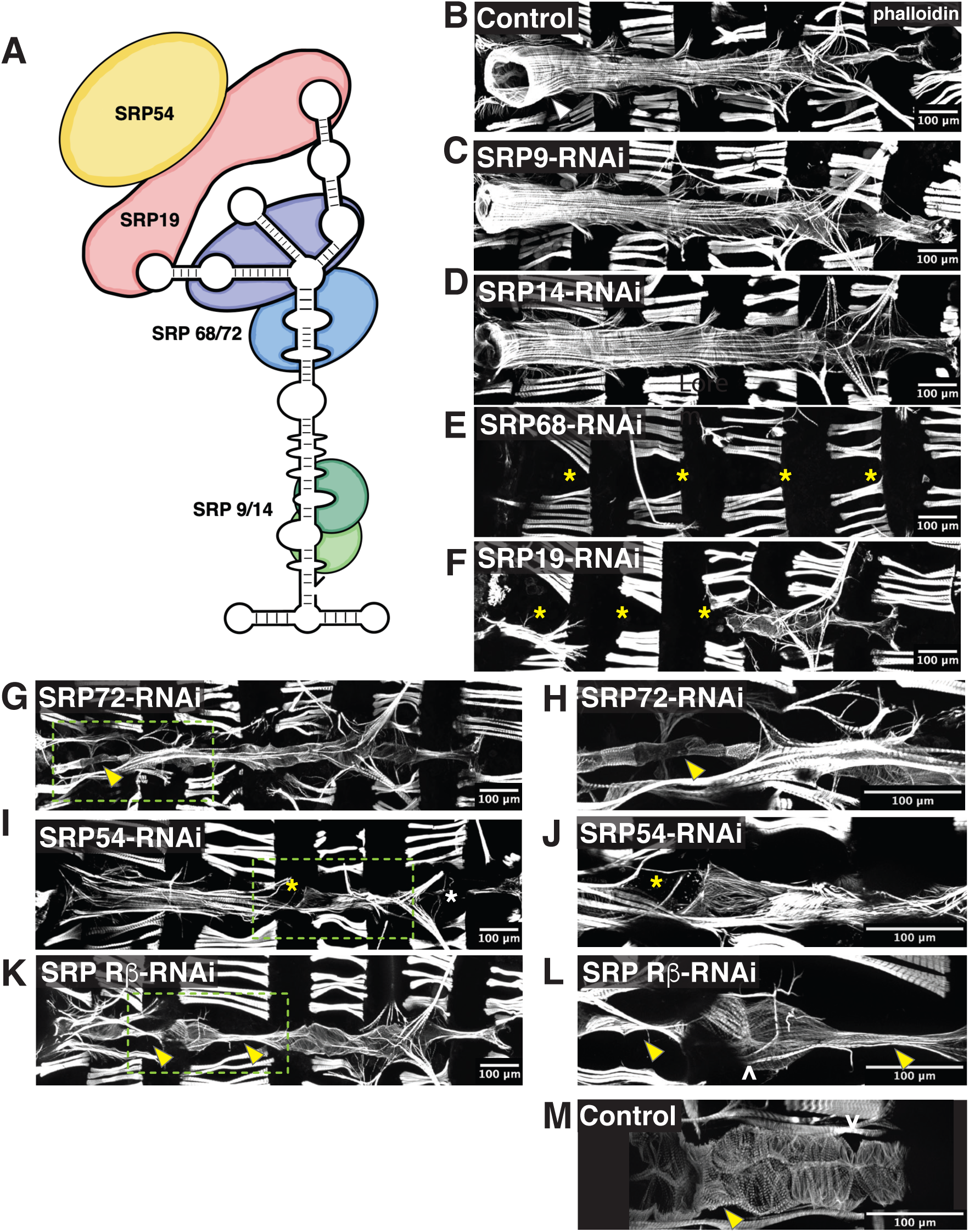
Knockdown (KD) of SRP subunits in the *Drosophila* heart caused distinct heart defects. **A**, The Signal Recognition Particle (SRP) is composed of an RNA molecule holding together 6 SRP subunits. **B-M,** Individual SRP subunits were KD by RNAi using the heart specific driver *Hand4.2*-GAL4 and adult flies were stained with phalloidin to determine their contribution to heart structure and development. * indicates missing heart segments or cardiomyocytes. KD of **C**, SRP9 and **D,** SRP14 subunits, responsible for elongation arrest during translation, did not lead to gross alterations in heart structure and were comparable to **B,** controls. **E**, KD of SRP68 led to complete loss of the heart. **G,** Interestingly, KD of SRP72, a binding partner to SRP68 led to the presence of a heart tube. **H,** Higher magnification of the conical chamber, marked by an arrowhead, showed that the conical chamber was constricted compared to controls, and resembled a larval aorta. **F**, KD of SRP19, led to a partial heart phenotype, where the anterior region of the heart was absent but the posterior end remained. **I**, KD of SRP54 led to a heart tube but with missing heart cells (indicated by asterisks) in random regions of the heart. **J**, Higher magnification of missing cardiomyocyte. **K**, KD of the SRP Receptor-*β*, a subunit of the receptor anchored to the ER membrane that binds to the SRP-ribosome-nascent chain complex, led to segments of the heart, usually the valves, that were constricted and larval like, indicated by the arrowhead. **L**, Higher magnification of the narrowed heart tube. Ostia structures are still present as marked by ^. **M,** As comparison, control valve cells in controls are wider than SRP*β* KD and are densely packed with myofibrils.

Each subunit of the SRP complex exhibits specialized roles in the binding and translocation process (**Figure 6A**) and therefore, each subunit could confer specialization of SRP function through recruitment of cofactors or selective targeting of nascent proteins. We therefore knocked down each of the SRP subunits individually, as well as the *β*-subunit of SRP Receptor (*SR-β*), to explore their role in heart development and how their phenotypes compare to *NACα* and *bic* KD. RNAi mediated KD of either *Srp9* or *Srp14* subunits in the heart driven by *Hand4.2*-GAL4 using two different RNAi lines did not produce gross differences in the heart structure compared to controls (**Figure 6 B-D**). In contrast, KD of *Srp68* led to complete absence of the adult heart, similar to *Nacα* KD (**Figure 6E**). Upon KD of *Srp72,* the heart tube was still present which is unlike the *Nacα* KD phenotype. The very anterior segment, called the conical chamber, remained very constricted, reminiscent of its larval structure (**Figure 6G,H**). This larval aorta-like structure suggests that this section of the heart failed to undergo remodeling. KD of *Srp19* led to a partial heart phenotype, where the anterior segment of the adult heart tube is absent, but retained a posterior segment, including the terminal chamber (**Figure 6F**), again unlike what was observed with *Nacα* KD. KD of *Srp54*, which recognizes and binds the signal sequence on the nascent polypeptide, led to a malformed heart with missing cardiomyocytes in random positions throughout the tube (**Figure 6 I,J**). KD of *SR-β* led to a heart tube with missing cardiomyocytes, which were most often the internal valves of the heart (**Figure 6K-M**). Interestingly, *in vivo* imaging of fluorescently labeled hearts in early pupae demonstrates that an intact larval heart tube forms with KD of any SRP components (**Supplemental Figure 6**), suggesting that their requirement prior to pupal remodeling of the heart may be less critical. Considering *Srp19* and *Srp68* KD largely abolished in adult hearts, the presence of an intact heart tube prior to cardiac remodeling in pupae suggests, that like *NACα*, their activity is critically required during cardiac remodeling and morphogenesis.

In summary, the KD of individual NAC and SRP subunits led to a range of distinct cardiac phenotypes that suggests each subunit and complex may function with distinct cell-type, regional and temporal specificities during cardiac development. Each subunit may contribute specialized function and interact with likely numerous cofactors that could target subsets of genes differently between SRP and *NACα*.

## Discussion

Nascent polypeptide Associated Complex (NAC) and the Signal Recognition Particle (SRP) are integral to protein biogenesis. Our work suggests that they are pertinent for establishing distinct proteomic landscapes that shape cell identity and direct developmental fates (**Figure 7**). We demonstrated a developmental role for the NAC subunit *Nacα*, in the *Drosophila* heart and Human Multipotent Cardiac Progenitors (MCPs) that influenced cardiac cell identity and morphogenesis through associations with *Hox* genes. These cardiac defects were triggered by knockdown (KD) of *Nacα* at specific stages in development. Similarly, disruptions in SRP subunit expression led to cardiac cell type specific defects in the fly and some changes in heart morphology occurred during later, critical stages of development. These results suggest that *Nacα* and SRP function and selection of protein targets were temporally regulated. Components involved in protein translation are a growing family of genes associated with tissue-specific and developmental defects [50], including the heart [51, 52]. For example, patients with Diamond Blackfan Anemia (DBA), a bone marrow failure syndrome caused by ribosomopathies, exhibit higher manifestations of CHD compared to the general population [53]. In other cases, ribosomopathies led to CHD defects without overt hematological abnormalities [9, 54, 55]. *Nacα* specifically has been associated with Tetralogy of Fallot [11] and increased myocardial mass [10]. Along the SRP pathway, *SRP54* mutations cause Schwachman-Diamond-like syndrome, a condition associated with CHDs [56, 57]. Furthermore, within the Pediatric Cardiac Genomic Consortium cohort [58], the SRP Receptor-*α* was associated with atrial septal defects while SRP Receptor-*β* was associated with cyanotic congenital heart disease including ventricular septal defect. As genomic studies of patients uncover new variants associated with CHD, a closer examination of variants within translational genes and their co-occurrence with gene variants within developmental genetic networks is warranted. The question remains how these generic proteins can achieve both tissue specificity and temporal control in their translational functions that are critical for normal cardiac cell identity and fly heart tube morphogenesis.

**Figure 7:**
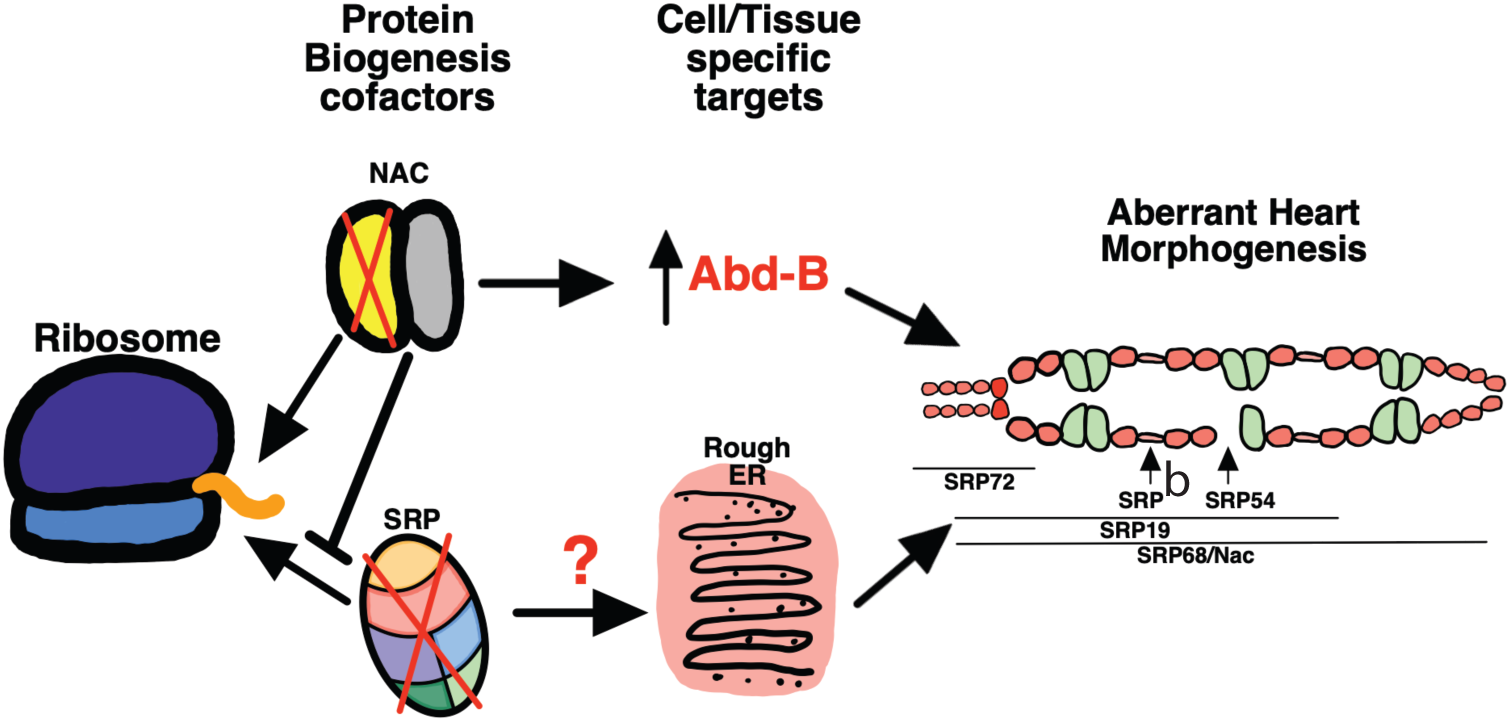
A schematic diagram of protein translation at the ribosome exit site, where nascent polypeptide cofactors NAC and SRP select for gene targets, regulating their expression in a tissue and/or cell specific manner to regulate heart morphogenesis. Our work suggests that the posterior determining *Hox* gene *Abd-B* is a target of *Nacα*, as the knockdown of *Nacα* led to *Abd-B* misexpression and disruption of heart morphogenesis. Knockdown of SRP subunits each led to distinct heart defects: SRP72 knockdown disrupts conical chamber morphogenesis, SRP-R*β* targets internal valve cells, SRP54 leads to missing cardiomyocytes, SRP19 leads to loss of anterior segment of the heart, while SRP68 led to a no heart phenotype similar to *Nacα* knockdown. Targets of SRP have yet to be determined.

Our studies found that KD of *Nacα* restricted by *Hand4.2*-GAL4 to mid- to late stages of embryo development (beginning stage 14 until hatching) specifically in cardiac relevant cells induced posterior terminal heart chamber defects that were evident in adult morphology (**Figure 4 E,F**). This suggests spatial specificity in *Nacα* function with embryonic origins targeting posterior cardiac cells more severely compared to more anterior regions. We also demonstrated that *Nacα* genetically interacts with a posteriorly expressed *Hox* gene, *Abd-B,* later during pupal stages, which further supports a stronger requirement for *Nacα* in the posterior heart; thus, heart defects occur from posterior to anterior regions with severity of KD, i.e. in embryo or pupae vs. both (**Figure 3**). These regional differences in *Nacα* activity is consistent with a previously described role for NAC in the posterior patterning of the developing *Drosophila* embryo [19, 20]. In the early syncytial embryo, posterior patterning proteins, such as OSKAR and NANOS, accumulate in the posterior end of the embryo. Components of the NAC complex were necessary to restrict translation and localization of OSKAR and NANOS to the posterior end [19, 20]. Absence of NAC components resulted in ectopic expression of posterior patterning proteins anteriorly leading to a lethal, bicaudal phenotype. How this translational control is achieved remains unclear, but could involve targeted translational pausing either through 3’UTR sequences or additional translation cofactors. In our work, the posterior preference is retained in an embryo where *Nacα* is knocked down only in the developing heart cells, which supports the idea that additional cofactors with posterior-anterior specific expression and/or function contribute to regulate *Nacα* activity.

Future studies will be aimed at determining the mechanisms that regulate *Nacα* activity that drives these anterior-posterior specific responses in the heart. Furthermore, how *Nacα* regulates *Hox* gene *Abd-B* repression in the anterior heart regions, and possibly other cardiac relevant targets, could help uncover pathways leading to targeted tissue defects and corresponding human diseases. Our data show that KD of *Nacα* is permitting *Abd-B* protein expression in the anterior regions of the larval heart and aorta, where it is normally absent (**Figure 3 C-E**). Ectopic *Abd-B* expression could occur through direct disinhibition of *Nacα* translational control of *Abd-B mRNA*. Alternatively, *Nacα* KD could have an indirect effect on cell identity that aberrantly turns on *Abd-B* transcription and subsequent protein expression. In addition, *Abd-B* protein is no longer strictly nuclear, rather is mistargeted and expressed throughout the cardiac cell of *Nacα* KD flies (**Figure 3E**). This could in part be a result of nascent protein mistargeting toward the SRP pathway that redirect *Abd-B* to the ER as well as secreted into the lumen. *Hox* genes play a crucial role in early cardiac specification, patterning and remodeling in the heart [59] and *Nacα* translational regulation could be a novel mechanism of *Hox* gene regulation that may contribute to pathogenesis of CHD.

Within the mammalian heart a similar role for regulated anterior-posterior *hox* gene expression is crucial for proper morphogenesis. Complex signaling events coordinate patterning of cardiac progenitor populations within the First and Second Heart Fields (SHF) [60]. Along the anterior and posterior axis of the SHF, cardiac progenitor cells are further subdivided into domains that express distinct transcriptional programs contributing to the development of distinct heart structures [61]. Characterization of the transcriptional profiles within subdomains have implicated specific *Hox* gene expression (*i.e. Hoxa1* and *Hoxb1*) to establish cellular identity and cell fate [61, 62]. Future work in mammals can help determine whether NAC and SRP activity may contribute to targeted expression of *Hox* genes within distinct subdomains of the SHF, which is suggested by our work in flies and human MCPs.

The effect of *Nacα* KD in fly hearts has a number of parallels to the targeted developmental mishaps resulting from mutation of another ubiquitous translational ribosomal protein, *Rpl38,* in mice. These mutant mice display specific axial skeletal patterning defects attributable to reduced association of a specific subset of *Hox mRNA* with polysomes (*Hox a4; a5; a9; a11; b3; b13; c8; d10),* which led to reduced protein levels of these *Hox* (validated for HOXA5, HOXA11 HOXB13 proteins), without changes in global protein translation [63, 64]. Mice with loss-of-function mutations in the targeted *Hox* genes (*Hoxa5, Hoxa9, Hoxa11, Hoxc8, Hoxd10)* phenocopy numerous aspects of the *Rpl38* mutant phenotypes along the rostro-caudal axis, suggesting that the altered translation of these *Hox* genes inflicted by *Rpl38* deficiency mediated the skeletal patterning defects. The results in flies and mice support the idea that seemingly generic components of the protein biogenesis pathway can impose a layer of translational control pertinent to tissue-specific development.

It is remarkable that the complete histolysis of the adult fly heart observed during pupal stages required KD of *Nacα* during both embryonic and pupal stages, either one alone was not sufficient (**Figure 4 D-H**). This was surprising, since during pupal stages the heart undergoes extensive remodeling, which requires significant protein biogenesis, and pupal-only KD of *Nacα* does not induce cell death, nor significant changes in heart structure or function. This suggests that *Nacα* function during embryonic stages may create a proteomic landscape establishing long-term programming of cardiomyocyte identity properly poised for future developmental responses, such as remodeling to the adult heart during metamorphosis. An embryonic requirement for later cardiac remodeling and survival was also found for ribosomal protein gene *RpL13* [9]. However, in the case for *RpL13,* KD in the embryo alone resulted in significant heart loss during pupal stages. Thus, again translational perturbances in the embryo can have long-lasting effects, suggesting a mechanism by which preprograming in the embryonic heart is critical for appropriate cardiac remodeling later in development. How translational proteins and mechanisms contribute to redefining cell identity would need to be better understood.

Our work using MCPs demonstrated a role for *Nacα* in human cardiac cell specification and differentiation. KD of *Nacα* redirected progenitor cell fates from a cardiomyocyte lineage toward a fibroblast lineage (**Figure 5**). In contrast, KD of cardiogenic transcription factors *Gata4/6* and *MyoCD*, directed progenitor cell fates toward a different population profile with decreased total cell population and decreased cardiomyocytes, but increased endothelial cells and no change in fibroblasts (**Figure 5**) [65, 66]. Although this difference in population profiles induced by KD of *Nacα* vs cardiogenic transcription factors suggests that *Nacα* utilizes distinct mechanisms for refinement of cardiac cell fates, it remains to be determined how these processes are coordinated.

Defects in differentiation and maturation caused by *Nacα* deficiency in flies are also observed in other cell lines and model systems including deviations in cell fates of osteoblast, myotubes and haematopoietic stem/progenitor cells [24, 26]. In higher order organisms, *Nacα* has evolved to encompass additional isoform-specific functions, including as a transcriptional coactivator [67]. Furthermore, a larger, alternatively spliced isoform of *Nacα* (*skNAC*), containing an additional proline-rich exon, is expressed specifically in striated muscle of vertebrates [68] and plays a role in transcriptional regulation, fiber-type specification, myofibrillogenesis, and sarcomerogenesis [27, 28, 31, 32, 68, 69]. This puts in question whether in flies *Nacα* only functions in translational regulation or may be involved in other cellular activities.

Protein biogenesis and its regulation at the ribosome exit site relies on a network of nascent protein chaperones. Because of the interactions between NAC and SRP that refine their ability to target specific nascent polypeptides emerging from ribosomes, we wanted to explore the effects of disrupting SRP activity on fly heart development and compare them to *Nacα* KD. We found that KD of individual SRP subunits led to unique cardiac phenotypes (**Figure 6**). While *Srp9* and *Srp14,* responsible for elongation arrest during translocation, did not alter adult heart structure, KD of *Srp68,* responsible for docking to the SR receptor on the ER membrane, led to complete loss of the heart. The subunit *Srp72* binds to *Srp68* and participates in SR docking activity, but its KD did not result in complete heart loss, but the defect was focused on the conical chamber of the heart, possibly hindering larval aorta transdifferentation into the adult fly heart during metamorphosis. KD of *Srp54*, a highly evolutionarily conserved protein subunit that bind to signal sequences on nascent polypeptides, led to missing cardiomyocytes at random positions in the heart. KD of *Srp19*, responsible for linking *Srp54* to the rest of the SRP complex led to a partial heart loss anteriorly, in contrast to *Nacα*,which preferentially deleted posterior portions of the heart. KD of the *β* subunit of the SR that binds nascent protein-SRP-ribosome complex at the ER membrane, led to dysmorphic cardiomyocytes, which were usually the internal valve regions. The varied phenotypes suggest specialized roles and a complexity in spatial regulation for each SRP subunit, similar to the posterior preference seen for *Nacα* KD. Furthermore, SRP’s influence on cardiogenesis is temporally regulated, as gross changes in morphology occur during crucial developmental stages, such as the disappearance of the heart during metamorphosis induced by *Srp68* KD (**Figure 6E**, **Supplemental Figure 6**).

Determining how KD of SRP subunits display cardiac tissue-specific effects would uncover novel aspects of protein biogenesis control in a cell-type specific manner. In contrast to *Nacα*, the SRP complex interacts with nascent proteins that are destined for insertion into the rough ER and subsequent translocation to the plasma membrane or for secretion. Therefore, SRP targets are likely enriched for membrane receptors. Thus, interruption of receptor function and signaling mediating non-cell autonomous cues may be partially involved in the cardiac phenotypes caused by disruption of SRP activity. For example, cardiac cells may respond aberrantly to ecdysone steroid hormone secreted by the ring gland during metamorphosis due to a lack of Ecdysone Receptor expression at the cardiac cell membrane. Indeed, this may in part be a contributing mechanism, as *Srp72* KD in the heart resulted in constriction of just the conical chamber in the anterior region of the adult heart, which resembled an unremodeled larval heart (**Figure 6G,H**). Still, why there appears to be regional specificity to this lack of Ecdysone Receptor response, despite *Srp72* KD throughout the heart, remains to be determined. Furthermore, KD of each SRP subunit resulted in differential effects on heart morphology that target discrete heart regions and cell types, suggesting a mechanism for tissue-specific regulation. It is possible that each SRP subunit binds a disparate set of differentially expressed cofactors to impart specific functional control of SRP activity.

In summary, our work demonstrated specific roles for the NAC and the SRP complex in cardiac development, using *Drosophila* and human MCPs, and offers novel mechanisms in the regulation of the cardiac proteome to establish cell identity and tissue patterning (**Figure 7**). How generic components of the translational machinery could lead to targeted effects on heart development and function is likely layered in a complex process that remains to be explored.

## METHODS

### Drosophila Strains

Heart-specific control of transcription was achieved by the GAL4-UAS system [70], in which the following cardiac relevant GAL4 drivers were used: Hand4.2-GAL4 [33, 71, 72], *tinCΔ*4-GAL4 [73], *tinD*-GAL4 [74], and *tinHE*-GAL4 [75]. *Dot-*GAL4 driver line [34] was used to express in pericardial cells. *Drosophila* GD and KK RNAi collection lines along with appropriate controls were obtained from the Vienna Drosophila Resource Center (VDRC)[76]. Two copies of a temperature-sensitive *tubulin*-GAL80 (*tubulin*-GAL80^ts^) were recombined and combined with the Hand4.2-GAL4 driver (Hand4.2-GAL4, Tubulin-GAL80^ts^; Tubulin-GAL80^ts^, HTT; [9]). All fly lines are listed in **Supplemental Table 1**.

### Developmental Regulation of Gene Expression

The HTT line allows for transcriptional control of constructs fused to a UAS enhancer by manipulation of ambient temperature. At 18°C, GAL80 is intact and blocks GAL4 mediated activation of the UAS enhancer. A shift to 29°C destabilizes GAL80 protein, permitting GAL4 binding to the UAS enhancer to induce transcription. As control, flies were placed in 18°C or 28°C throughout development until eclosion when an intact or an absent heart were expected, respectively. For *Nacα* knockdown (KD) in embryonic stages, female and male flies were placed in 29°C and embryos collected every 2 hours. Embryos were maintained in 29°C for 24 hours and then shifted to 18°C for the remainder of development. Another set of embryos were maintained in 29°C for 48 hours before shifting to 18°C. For *Nacα* KD during pupal stages, fly crosses were maintained in 18°C until wandering 3^rd^ instar larvae were collected. They were then immediately placed in 29°C until eclosion. For KD of *Nacα* starting at mid-larvae, embryos from fly crosses at 18°C were collected every 2 hours, maintained in 18°C for 16 days when larvae were approximately at L2 stages. Larvae were then placed in 29°C until eclosion. Finally, to test *Nacα* KD during embryonic and pupal stages only, embryos were collected from fly crosses at 29°C for 24 hours as described above, and then returned to 18°C to develop through larval stages. Wandering larvae were then collected and then placed in 29°C until eclosion. Adult flies were assessed for heart function using SOHA (see below) and immunostained to examine structure (see **Figure 4**).

### Immunostaining of Adult Drosophila Hearts

Adult flies were dissected and treated with 10mM EGTA in PBT (PBS + Triton-X-100; 0.03% Triton X-100), for 2 minutes to maintain a relaxed state of the heart. Hearts were then fixed with 4% PFA in PBT for 20 minutes, followed by three 10-minute PBT washes. Hearts were stained with primary antibodies (EC11 *Pericardin*, Developmental Studies Hybridoma Bank, DSHB) and incubated overnight in 4°C. Hearts were then washed with PBT three times for 15 minutes each, followed by incubation with fluorescent secondary antibodies (1:500, Jackson ImmunoResearch Laboratories, Inc.) and Alexa Fluor conjugated phalloidin (1:300, Life Technologies) in 4°C overnight. Hearts were then washed with PBT three times for 15 minutes each and then once with PBS. Hearts were mounted using ProLong Gold Mountant with DAPI (Life Technologies). Immunostained preparations were visualized with an Imager.Z1 equipped with an Apotome2 (Carl Zeiss, Jena), Hammamatsu Orca Flash4.0 camera, and ZEN imaging software (Carl Zeiss).

### Live-imaging of the Heart in Drosophila Pupae

For fluorescence-based heart function analysis, we crossed a fly line that contained both Hand4.2-GAL4 and a heart enhancer fused to tandem-Tomato fluorescent protein (tdtK), to controls, *Nacα*-RNAi, or *bicaudal-RNAi* flies [76]. White pupae (WP) from these crosses were collected and lined up on small petridishes with dorsal side facing up. A subset of WP was imaged immediately while another subset was aged for approximately 20 hours in room temperature. Dishes were placed on a Zeiss Imager M1 equipped with a Hamamatsu Orca-Flash 4.0 Digital Camera (C11440), and heart images were captured every 2 minutes, for a duration of about 55 hours at room temperature (approx. 20°C) using ZEN imaging software (Carl Zeiss). Movies were formatted and compiled using ImageJ and iMovie (Apple).

### Heart Function Analysis

Assessment of *Drosophila* heart function and structure using the Semi-automatic Optical Heartbeat Analysis (SOHA) method as previously described [77, 78]. Briefly, four-day old adult flies were anaesthetized with FlyNap (Carolina Biological Supply Co, Burlington, NC) and dissected in oxygenated artificial hemolymph to expose the beating heart within the abdomen. Hearts were placed on an Olympus BX61WI microscope while being filmed through a 10x water immersion lens with a high-speed digital camera (Hamamatsu Photonics C9300 digital camera) using HCI image capture software (Hamamatsu). High-speed movies were analyzed using the Semi-automated Optical Heartbeat Analysis (SOHA) software [77, 78]. Parameters measured include Heart Period (HP), Diastolic Interval (DI), Systolic Interval (SI), Arrhythmicity Index (AI), Diastolic Diameter (DD), Systolic Diameter (SD) and Fractional Shortening (FS) (**Figure 2A**). FS, a measure of contractility, is calculated using the following equation FS=(DD-SD)/DD.

### MCP cell culture, siRNA transfection, and Immunostaining

A pool of 4 unique siRNA sequences targeting different regions of selected genes and random control were obtained from the human siGENOME library from Dharmacon, Inc. Frozen 5-day old human MCPS [44, 79] were thawed and transfected with siRNAs at 5nM final concentration using Lipofectamine RNAiMAX transfection reagent (Invitrogen). Approximately 20,000 cells were plated in each well of a 384-well plate (Greiner Bio-One) coated with Matrigel Basement Membrane Matrix (Corning). Cells were incubated at 37°C and media refreshed every second day. Each experiment contained quadruplicate technical replicates per condition and performed on different batches of MCP clones programmed independently.

Immunostaining to identify cell-type composition of cultures were performed 9 days following siRNA transfection. Cells were fixed with warmed 4% paraformaldehyde solution for 30 minutes without agitation, followed by an additional 30 minutes of fixing with agitation. Wells were washed with Phosphate Buffered Saline (PBS) for 10 mins and repeated three times. Samples were then incubated with blocking solution for 30 mins (10% horse serum, 2.5% Triton-X-100, 10% gelatin). Primary antibodies were diluted in blocking solution and added to samples for 1 hour at room temperature (RT): ACTN1 (Sigma, A7811), TAGLN (Abcam, Ab14106), CDH5 (R&D Systems, AF938). Wells were washed with PBS for 15 mins and repeated three times. Samples were incubated with Alexa-conjugated secondary antibodies (Life Technologies) including DAPI diluted in blocking solution for 1 hour at RT. Wells were washed for 15 minutes and repeated three times, following which samples were imaged using a High-Throughput microscope (ImageXpress, Molecular Devices). Fluorescence was quantified using custom MetaXpress software (Molecular Devices) whereby, the number of total cells and cells positive for ACTN1, TAGLN, and CDH5 in each sample were quantified [44, 79].

### Statistical Analysis

Statistical analysis was performed using GraphPad Prism (GraphPad Software, La Jolla USA). Data are presented as the mean ± SEM. Human MCP, cardiomyocyte and fibroblast data were analyzed using a one-way ANOVA analysis followed by a Dunnett’s multiple comparison post-hoc test. A student’s unpaired t-test was used to analyze *Drosophila* heart function data.

## SUPPLEMENTAL FIGURE LEGENDS

**SUPPLEMENTAL FIGURE 1:**
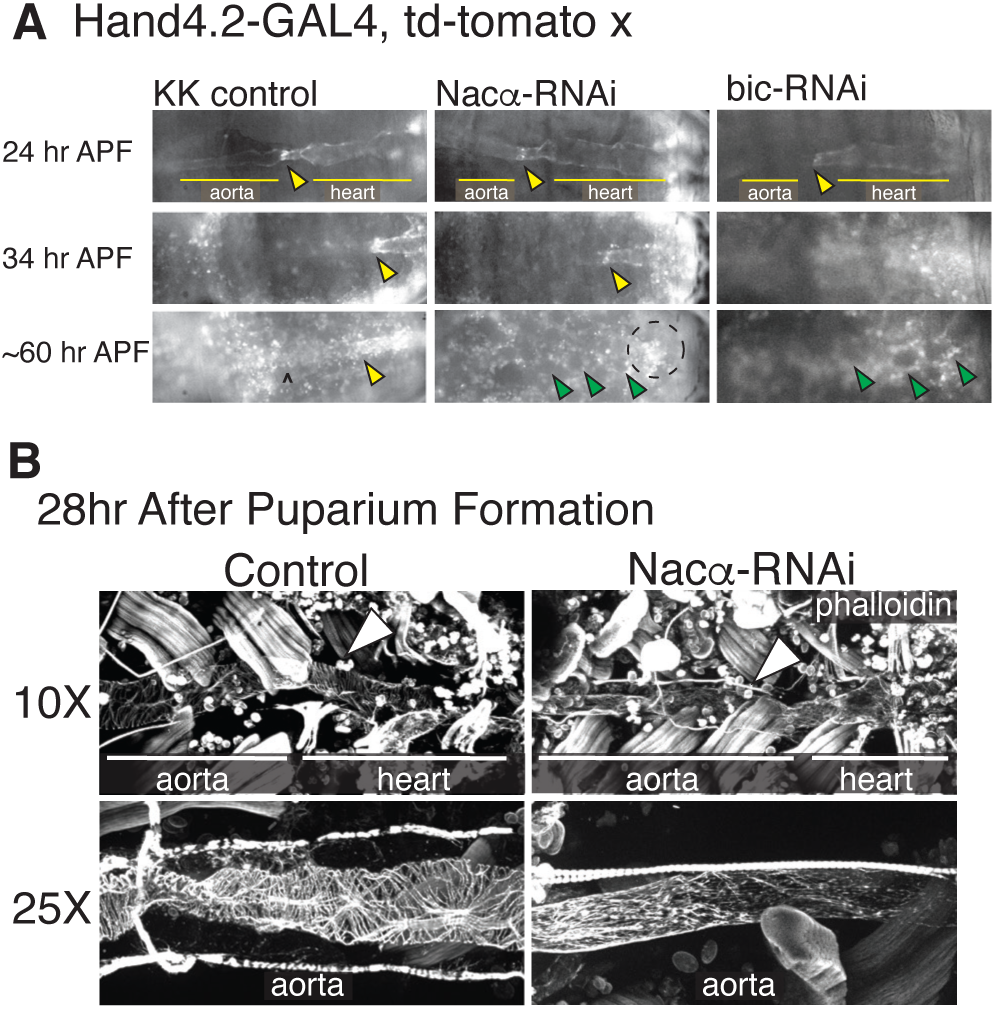
**A,** Still images of remodeling pupal hearts with control, *Nacα* knockdown (KD) and *bic* KD. At 24hr APF, the early pupal heart is present in controls (left) and *Nacα* (middle) and *bic* (right) knockdown flies. The internal valves (yellow arrowheads) are visible which separate the larval aorta (left) and the heart (right) which we use as a landmark through remodeling. At about 34hr APF, the internal valves are present but the aorta is more difficult to visualize as the heart transdifferentiates. At approximately 60hr APF, the remodeling adult heart tube is visible in controls with identifiable ostia structures (marked by ^). In *Nacα* and *bic* KD hearts, no heart tube is visible and the area is filled in by rounded fat cells (green arrowheads). The area of the embryo with fluorescent signal (circled) are remnants of histolyzed cardiomyocytes that slowly disperse and weaken in intensity. **B,** Pupal dissections at approximately 24-26hr APF stained with phalloidin shows the presence of the fly aorta and heart in controls (left) and with *Nacα* KD (right) at two magnifications. Arrows point to the presence of a heart tube.

**SUPPLEMENTAL FIGURE 2:**
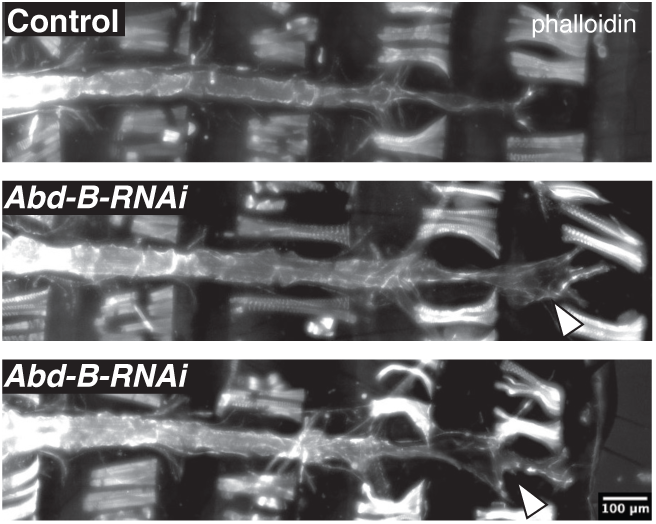
Knockdown of *Abd-B* using heart specific driver *Hand4.2*-GAL4 led to intact hearts with posterior ends (indicated by arrowhead) that were more prominent and dilated compared to controls suggesting incomplete histolysis during cardiac remodeling.

**SUPPLEMENTAL FIGURE 3:**
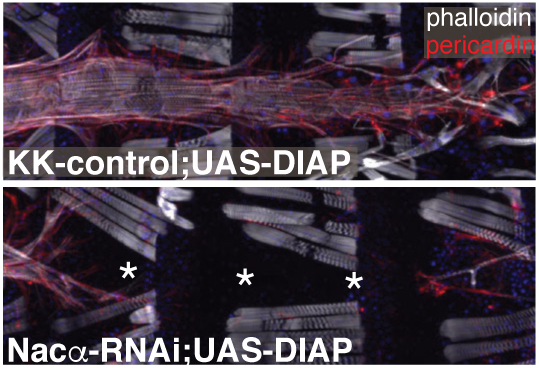
Overexpression of an inhibitor of apoptosis (Diap1) concurrently with *Nacα-RNAi* does not rescue the loss of the heart. * indicates absence of the adult heart tube.

**SUPPLEMENTAL FIGURE 4:**
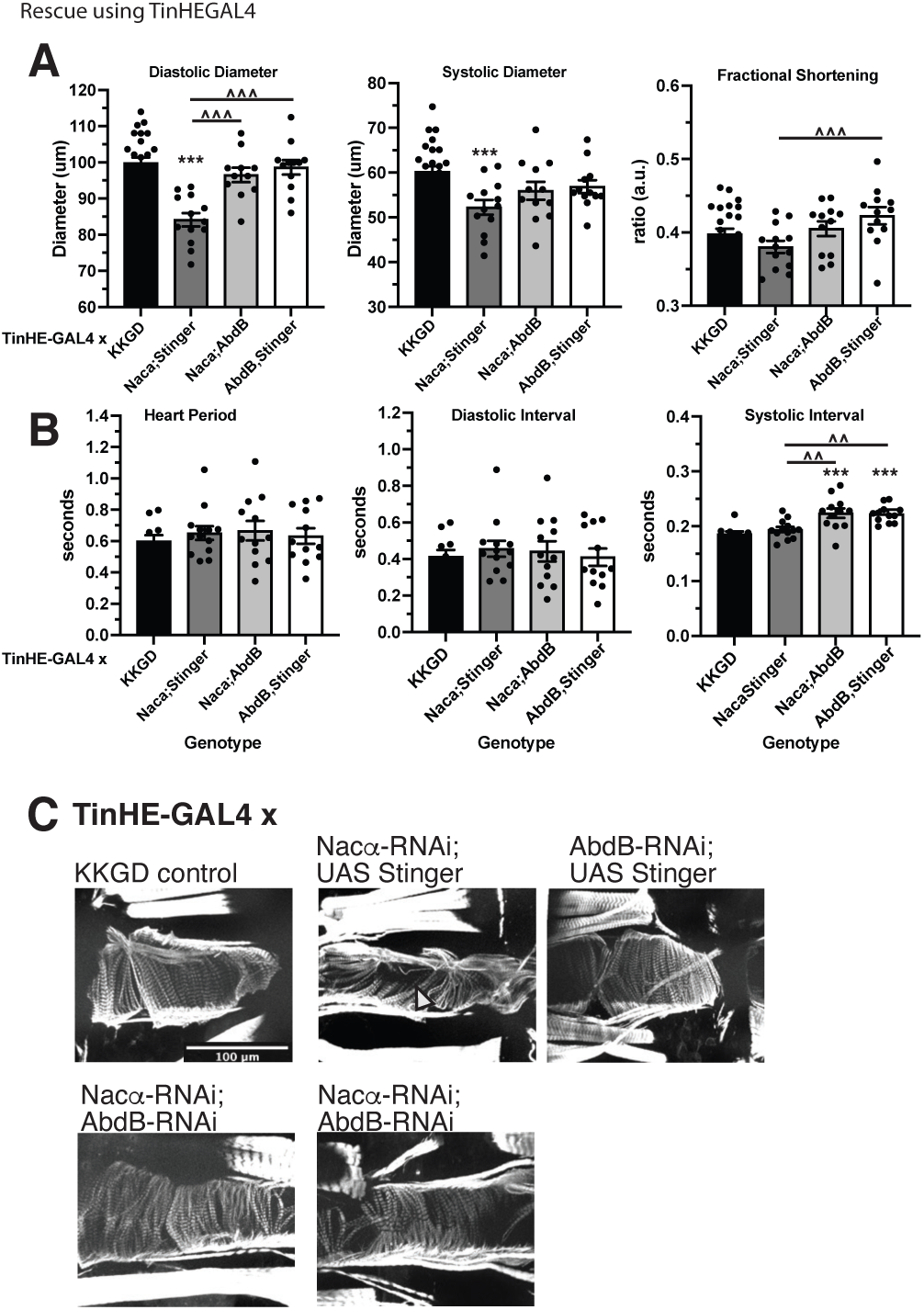
Testing interaction of *Nacα* and *Hox* gene *Abd-B* using the weaker cardiac specific *tinHE*-GAL4 driver by **A,** functional, **B,** temporal and **C** structural assessment. **A,** Knockdown (KD) of *Nacα* (combined with UAS-Stinger::GFP to control for UAS binding sites) using *tinHE*-GAL4 caused a decrease in both diastolic and systolic diameters that produced a slight but not significant decrease in fractional shortening. KD of *Abd-B* (combined with UAS-Stinger::GFP) did not produce significant changes in fractional shortening or diameters compared to control but fractional shortening and diastolic diameters were significantly higher compared to *Nacα;Stinger* genotype. Combined knockdown of *Nacα* and *Abd-B* produced heart parameters that were not different to controls but recapitulated heart function produced by *Abd-B* KD alone, suggesting that the heart function was rescued. **B,** Temporal parameters were unchanged with *Nacα*-RNAi expression. KD of *Abd-B* lengthened systolic interval compared to controls. Combined *Nacα* and *Abd-B* KD displayed longer systolic intervals similar to *Abd-B* KD alone, suggesting a rescue. **C,** Phalloidin staining of select genotypes. Compared to controls, *Nacα* knockdown disrupted circumferential fiber organization creating gaps in the matrix (similar to Figure 2I). KD of *Abd-B* did not significantly alter circumferential fiber organization. Combined knockdown of *Nacα* and *Abd-B* (2 examples shown) improved circumferential fiber organization compared to *Nacα* knockdown alone. * vs control KKGD. ^ compared to *Nacα*;*Stinger.* * *p<0.05, ** p<0.01, *** p<0.001*.

**SUPPLEMENTAL FIGURE 5:**
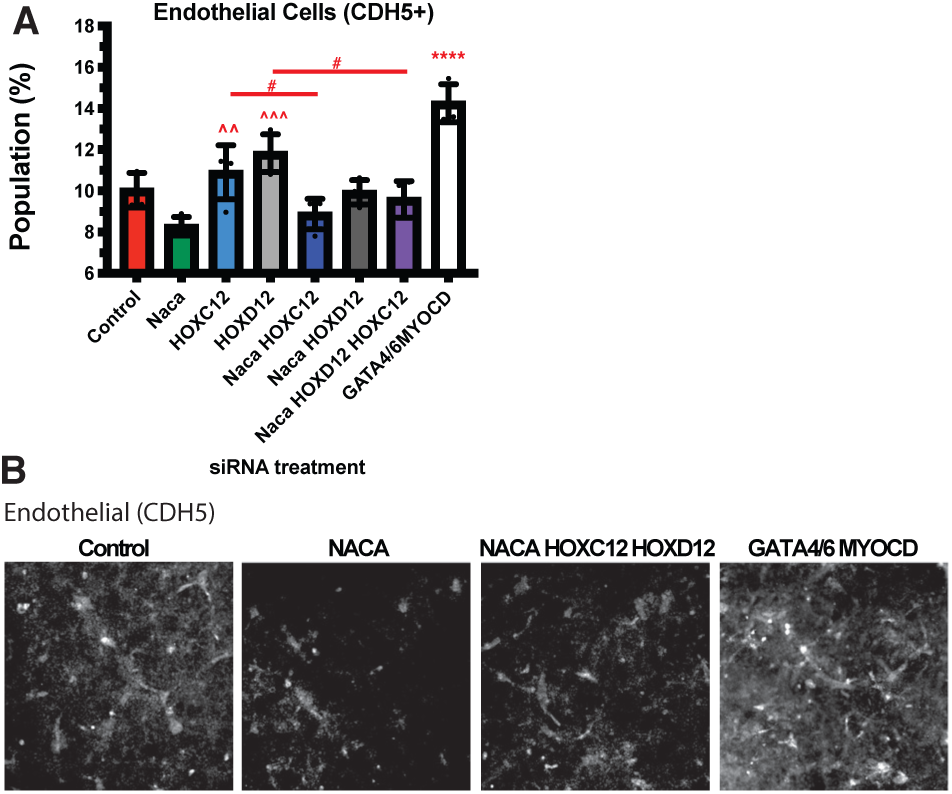
*Nacα* and Hox genes interact to redirect differentiation of Multipotent Cardiac Progenitors (MCPs). **A,** Knockdown (KD) of *Nacα,* Hox genes or their combination did not produce a significant change in the proportion of endothelial cell (CDH5+). The KD of transcription factors *Gata4/6,MyoCD* increased the proportion of endothelial cells. **B,** Representative images of immunohistological staining for select conditions.

**SUPPLEMENTAL FIGURE 6:**
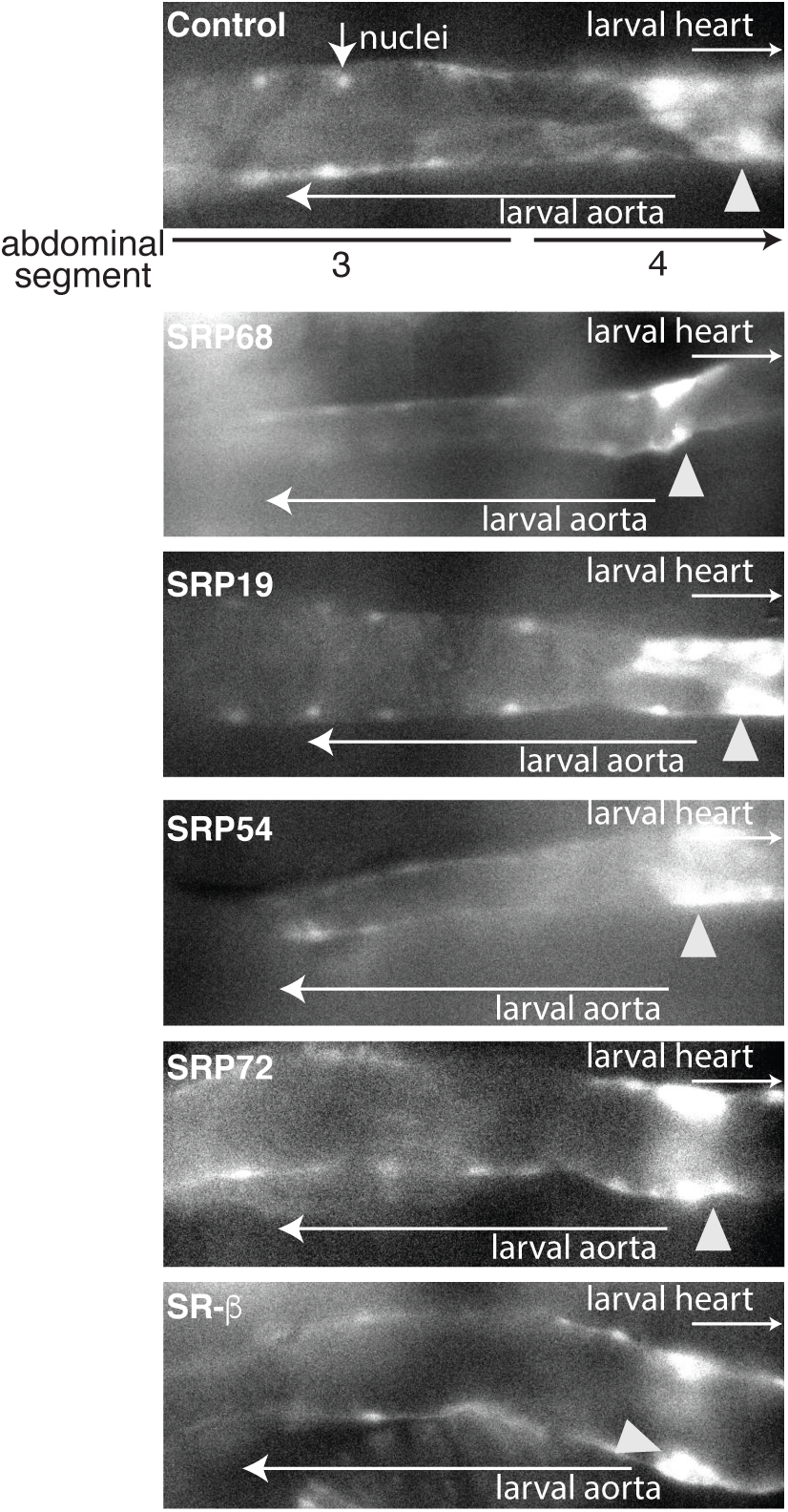
Images of fluorescently labeled (tdtK) larval aorta and heart (abdominal segments 3 and 4) of White Pupae, which showed that heart tubes were present prior to cardiac remodeling with any of the SRP subunit knocked down using *Hand4.2*-GAL4. Internal valves separating the larval aorta and heart are marked by arrowheads.

**SUPPLEMENTAL VIDEO 1, 2 and 3**: Video of fluorescently labeled pupal hearts (tdtK) of control (**Video 1**), *Nacα* (**Video 2**) and *bicaudal* (**Video 3**) knockdown flies. Images were captured starting early pupae (about 6-7 APF) through cardiac remodeling up until about 80 hr APF.

**SUPPLEMENTAL TABLE 1:** Table of transgenic lines used in the study.

| Fly gene | Human Gene | DIOPT score | Fly Collection | ID/Stock |
| --- | --- | --- | --- | --- |
| *Nac-alpha* | *NACA* | 11 | VDRC | KK v109114<br>GD v36017 |
| *bicaudal* | *BTF3/NACβ* | 12 | VDRC | KK v104718<br>GD v15453 |
| *AbdB* | *HOXC12*<br>*HOXD12*<br>(multiple orthologs) | 3<br>3 | VDRC | KK v104872<br>GD v12024 |
| *UAS-AbdB* |  |  | BDSC | 913 |
| *SRP9* | *SRP9* | 13 | VDRC | KK v105682<br>GD v23678 |
| *SRP14* | *SRP14* | 11 | VDRC | KK v101444<br>GD v23422 |
| *SRP19* | *SRP19* | 14 | VDRC | KK v106756 |
| *SRP68* | *SRP68* | 14 | VDRC | KK v104867<br>GD v273351 |
| *SRP72* | *SRP72* | 14 | VDRC | GD v43978<br>GD v21641 |
| *SRP54* | *SRP54* | 11 | VDRC | GD v51088 |
| *UAS-stinger* | *nuclear GFP* |  | BDSC | 84277 |
| *UAS-Val10-GFP* |  |  | BDSC | 35786 |
| *tubulin-GAL80ts* | *Tubulin* |  | BDSC<br>BDSC | 7017<br>7019 |
| *Hand4.2-GAL4*<br>*Hand4.2-GAL4; tdtK* | *Hand* |  |  |  |
| *tinHE-GAL4* | *tinman* |  |  |  |
| *tinCD4-GAL4* | *tinman* |  |  |  |
| *Dot-GAL4* | *UDP glucuronosyltransferase family* |  |  |  |

